# A highly conserved neuronal microexon in DAAM1 controls actin dynamics, RHOA/ROCK signaling, and memory formation

**DOI:** 10.1101/2023.01.12.523772

**Authors:** Patryk Poliński, Marta Miret Cuesta, Alfonsa Zamora-Moratalla, Federica Mantica, Gerard Cantero-Recasens, Davide Normanno, Luis P. Iñiguez, Cruz Morenilla-Palao, Patricia Ordoño, Sophie Bonnal, Jonathan D Ellis, Raúl Gómez Riera, María Martínez De Lagrán Cabredo, Álvaro Fernández-Blanco, Cristina Rodríguez-Marin, Jon Permanyer, Orsolya Fölsz, Eduardo Dominguez-Sala, Cesar Sierra, Diana Legutko, José Wojnacki, Juan Luis Musoles Lleo, Francisco José Muñoz López, Benjamin J Blencowe, Eloisa Herrera, Mara Dierssen, Manuel Irimia

## Abstract

Actin cytoskeleton dynamics is critical for nervous system development and function, yet the role of alternative splicing in controlling these processes is poorly understood. A highly conserved subset of neuronal-specific microexons coordinates fundamental aspects of nervous system biology. A subset of these exons is enriched in genes involved in actin cytoskeleton, yet their functions are unknown. Here, we focus on a microexon in DAAM1, a member of the formin-homology-2 (FH2) domain class of proteins, which have diverse functions associated with the reorganization of the actin cytoskeleton. Remarkably, splicing of the microexon extends the linker region of the DAAM1 FH2 domain and leads to qualitative and quantitative changes in actin polymerization. Deletion of the microexon results in neuritogenesis defects and increased calcium influx in differentiated neurons. Moreover, mice harboring the deletion exhibit postsynaptic defects, reduced number of immature dendritic spines, impaired long-term potentiation, and deficits in memory formation. These deficits are associated with increased RHOA/ROCK signaling, pivotal in controlling actin-cytoskeleton dynamics, and were rescued by treatment with a ROCK inhibitor. We thus demonstrate that a conserved neuronal microexon in DAAM1 is critical for controlling actin dynamics through the RHOA/ROCK signaling pathway and is necessary for normal cognitive functioning.

## Introduction

Higher cognitive functions of mammalian brains result from complex interactions among billions of neurons mediated by synapses. These connections require the formation of specialized cellular structures in the presynapse, responsible for the proper storage and turnover of synaptic vesicles filled with neurotransmitters, and in the postsynapse, which receives these signals mainly in the dendritic spines. The precise organization of both the pre- and post-synaptic terminals is, to a great extent, dependent on the actin cytoskeleton (Cingolani and Goda 2008). At the presynapse, actin networks are responsible for the spatial segregation and cycling of synaptic vesicles (Dillon and Goda 2005; Papandréou and Leterrier 2018), while in the postsynaptic terminals, they drive spine morphology and neurotransmitter receptor mobility (Cingolani and Goda 2008). As a result, the actin cytoskeleton is crucial for multiple brain functions, including memory formation. For example, actin dynamics directly affects experience-dependent synaptic plasticity, which induces specific connectivity patterns between the synapses (Lamprecht 2021; McLeod and Salinas 2018).

In recent years the Rho family of small GTPases have been described as the key regulators of synaptic plasticity due to their control over various molecules essential for actin cytoskeleton assembly. RHOA is one of such GTPases, as it controls immature dendritic spine number by regulating the activity of the ROCK signaling pathway (Zhang et al. 2021). Moreover, RHOA directly interacts with formins, a group of actin nucleating proteins that are critical drivers of actin dynamics (Mattila and Lappalainen 2008; Pollard 2016). Formin proteins are remarkably diverse and versatile, with 15 members grouped into seven families in humans (Schönichen and Geyer 2010). All of them are characterized by a conserved Formin-homology-2 (FH2) domain that forms homodimeric structures directly responsible for the processive polymerization of F-actin filaments (Kühn and Geyer 2014; Yamashita et al. 2007; Gao and Chen 2010). The FH2 domain-based “tethered dimer” exists at two states in equilibrium that allow either actin-binding or dissociation. Notably, the interconversion between these two states occurs through the dynamic expansion of the FH2 ring and is possible thanks to the high flexibility provided by the linker region (Schönichen and Geyer 2010; Xu et al., 2004; Yamashita et al. 2007).

Alternative splicing, the differential processing of exons and introns is a widespread mechanism contributing to the diversification and specialization of protein function. Most human multi-exonic genes undergo alternative splicing (Pan et al. 2008; E. T. Wang et al. 2008), and its prevalence is exceptionally high in mammalian brains (Barbosa-Morais et al. 2012). Neural-specific alternative splicing is relatively conserved throughout vertebrate evolution (Barbosa-Morais et al. 2012), and its misregulation has been linked to neurodevelopmental disorders (Feng and Xie 2013). A particularly striking example is provided by microexons, a class of 3-30 nt-long exons with a high degree of neuronal-specific inclusion and evolutionary conservation, and whose skipping has been linked to autism spectrum disorder and cognitive dysfunction in human and animal model studies (Irimia et al. 2014; Quesnel-Vallières et al. 2016; Parras et al. 2018; Gonatopoulos-Pournatzis et al. 2018; 2020; Torres-Méndez et al., 2022). Of several microexons that have been functionally studied, critical roles have emerged in processes including gene regulation, neuritogenesis, formation of synaptic connections, and behavior (Gonatopoulos-Pournatzis and Blencowe 2020; Dai, Aoto, and Südhof 2019; S. Wang et al. 2024). However, most microexons have not been functionally characterized on any level. In particular, genes harboring neuronal microexons are highly enriched in functions related to actin filament organization, actin filament-based processes, and cytoskeletal protein binding (Irimia et al. 2014; Quesnel-Vallières et al. 2016), suggesting important roles in these processes. Nonetheless, the impact of microexon inclusion on actin-related neuronal processes remains unknown.

In this study, we investigate the function of a neuronal-specific microexon in the Dishevelled-associated activator of morphogenesis 1 (DAAM1) and show that it impacts its FH2 domain, modifying the length of its linker region. Using biochemical assays and TIRF microscopy we demonstrate that splicing of this microexon modulates actin polymerization dynamics and fiber structure. Moreover, microexon inclusion is essential for proper neuronal function, since its deletion in differentiated in vitro results in increased Ca2+ influx upon depolarization, and increased activity of the RhoA/ROCK signaling pathway. Consistent with these results, deletion of the microexon *in vivo* results in multiple abnormalities in mice, including a decreased number of thin dendritic spines, reduced long-term potentiation (LTP), and deficient memory formation. Remarkably, treatment with an RHOA/ROCK signaling cascade inhibitor rescues phenotypes resulting from the deletion of the microexon, including altered transcriptomic signatures in differentiated neurons and memory impairments *in vivo*.

## Results

### DAAM1 harbors a highly conserved neural-specific microexon within its FH2 domain

To investigate the contribution of alternative splicing to the functional specialization of actin polymerization and nucleation in mammalian neurons, we focused on the family of formins, comprising 15 genes in mammals. Using data from VastDB (Tapial et al. 2017), we found that these genes were widely expressed across cell and tissue types, and eleven of them showed substantial levels of expression (TPM ≥ 10) in brain samples (Figure 1a). Moreover, seven increased their expression during neuronal differentiation, with the Daam1 showing the highest absolute expression in neurons (Figure 1a, right panel). Regarding alternative splicing, we identified six exons in four formin genes with different degrees of neural-specific splicing regulation (Figures 1b,c; see Methods) and evolutionary conservation across vertebrates (Figure 1d). At the protein level, one of these exons was predicted to disrupt the open reading frame upon inclusion (Formine Like 3, FMNL3), two were second-to-last exons containing stop codons giving rise to alternative C termini (FMNL1-B and FMNL2-B), and three corresponded to microexons that preserve the reading frame (DAAM1, FMNL1-A and FMNL2-A)(Figure 1b). As generally described for microexons (Irimia et al. 2014), these three exons fell either within a structured domain (i.e., the FH2 domain in DAAM1) or right next to it (i.e., the GTPase-binding domain in FMNL1 and the FH2 domain in FMNL2) (Figure 1b).

**Figure 1.**
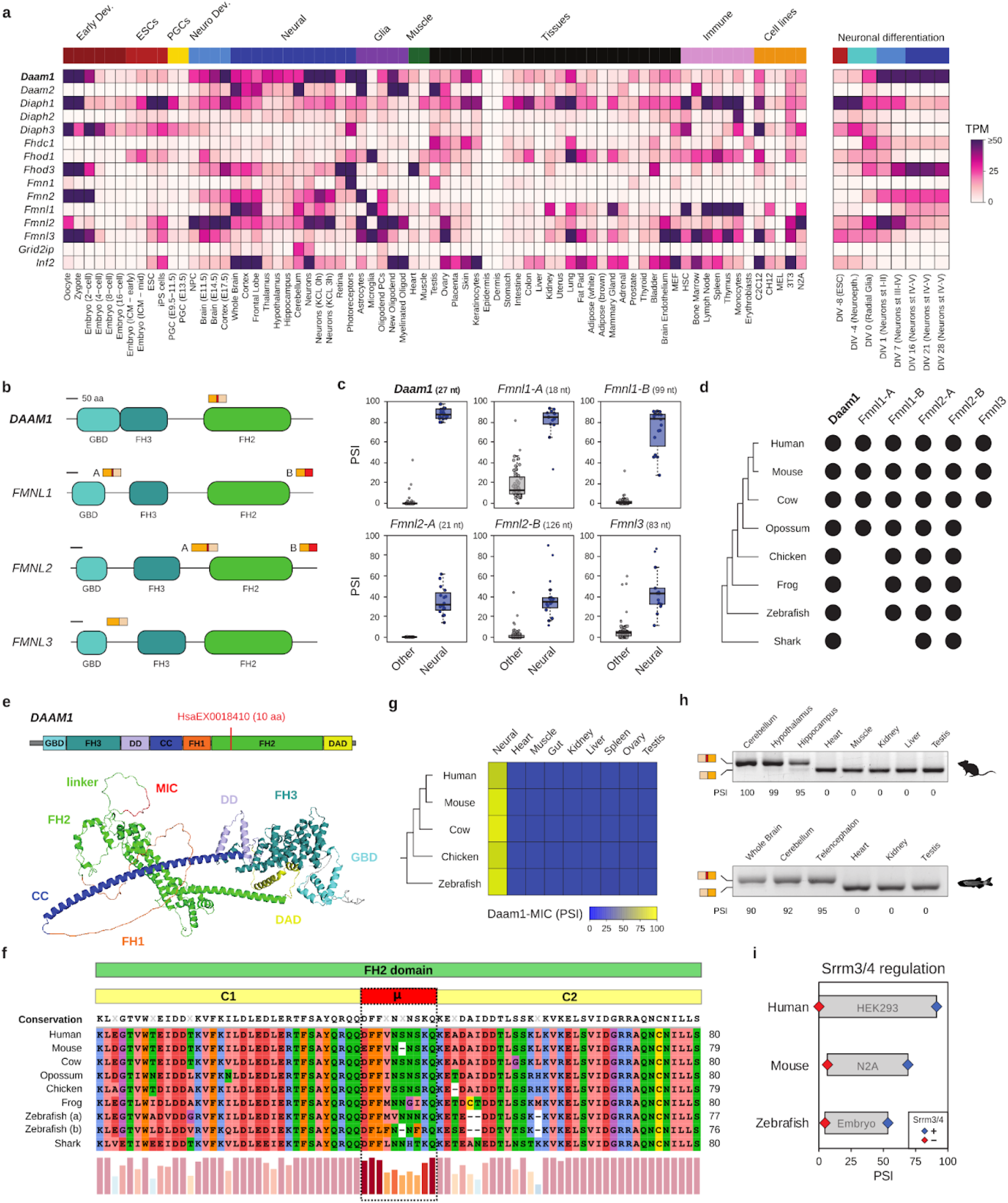
Neuronal-specificity anti evolutionary conservation of Daaml-MIC. (a) Heatmap showing gene expression levels of formin genes across multiple tissues and cell types based on VastDB. TPM: Transcript Per Million, (b) Schematic representation of the protein impact of neural-specific exons. Yellow-colored boxes represent upstream and downstream exons where the red box represents the alternatively spliced exon. GBD, GTPase-binding domain; FIB, Formin-Homology-3 domain or Diaphanous-Inhibitory Domain; FH2, Formin-Homology-2 domain, (c) Distribution of inclusion levels, using the Percent Spliced In (PSI) metric, for neural-specific exons in formin genes. PSI values were obtained from VastDB. (d) Evolutionary conservation of neural-specific alternative exons in formin genes. Black dots indicate the presence of an exon ortholog at the genome level, (e) Schematic representation of DAAM1 and its domains (top), together with DAAMI protein structure based on AlphaFold2 (bottom). The location of Daaml-MIC (HsaEX00l8410 in VastDB) is shown. GBD, GTPase-binding domain; FH3, Formin-Homology-3 domain or Diaphanous-Inhibitory Domain; DD. Dimerization Domain; CC, Coiled-Coil Domain; FH1, Formin-Homology-1 domain; FH2, Formin-Homology-2 domain; DAD, Diaphanous-Autoregulatory Domain, (f) Partial amino acid sequence alignment of the FH2 domain of DAAM 1 orthologs across vertebrates. Microexon (p), upstream (Cl), and downstream exons (C2) are indicated. The barplot depicts amino acid conservation, (g) Conserved neural-specificity of Daaml-MIC orthologs in vertebrates. PSI values from VastDB. (h) RT-PCR assays showing the inclusion of Daaml-MIC orthologs in different tissues from mice and zebrafish. Inclusion and the skipping bands are indicated on the left side of the gel. PSI values are indicated below, (i) Srrm3/4-dependent regulation of Daaml-MIC orthologs in human, mouse and zebrafish. PSI values in the condition with (+, blue) or without (-, red) Srrm3/4 are shown. RNA-seq data from human HEK293 cells overexpressing human SRRM4 (Torres-Mendez et al. 2019), mouse N2A cells upon Srrm3/4 knockdown (Gonatopoulos-Poumatzis et al. 2018), and zebrafish retinae extracted from Srrm3 KO 5 days post fertilization larvae (Ciampi et al. 2022).

In particular, the microexon in DAAM1 lies within the linker region of the FH2 domain and modifies its length (Figure 1e), which has been reported to have a significant impact on actin polymerization (Yamashita et al. 2007). Protein alignments of the FH2 domain of DAAM1 and its linker region across vertebrates revealed that the microexon has been conserved at the orthologous position from sharks to humans, keeping a constant length of 9-10 amino acids (Figure 1f). Moreover, transcriptomic data from various vertebrate species available in VastDB (Tapial et al. 2017) showed that the inclusion of the microexon in DAAM1 is also tightly restricted to neural tissues in other vertebrates (Figure 1g). This inclusion pattern was validated using RT-PCR assays in mouse and zebrafish tissues (Figure 1h). Additionally, in line with previous studies (Calarco et al. 2009; Raj et al. 2014), we found that the inclusion of the microexon was highly dependent on the action of the neuronal-specific Ser/Arg Repetitive Matrix 3 and 4 genes (SRRM3/4) in humans, mice and zebrafish (Figure 1i). Because of its higher neural-specificity and evolutionary conservation compared to other formin alternative exons and its direct impact on the FH2 domain, we decided to investigate the biological relevance of the microexon of DAAM1 (hereafter Daam1-MIC) at the molecular, cellular, and organismal levels.

### Daam1-MIC extends the FH2 linker region, impacting actin polymerization and structure

To gain further insights into the functional impact of Daam1-MIC, we generated models of the core structure of DAAM1’s FH2 domain (Figure 2a) for the inclusion and skipping variants. These models suggest that the microexon is inserted into the disordered linker region of FH2 without causing major structural changes (red, Figure 2b). This linker region is responsible for both the flexibility of the FH2 domain and its processive polymerization of actin (Otomo et al. 2005). In addition, a comparative analysis among human formin proteins revealed that the insertion of the ten amino acids encoded by the microexon makes the linker of DAAM1 the longest among all formins (Figures 2c,d, and S1a). Since artificial versions of the linker with varying lengths were previously associated with DAAM1 activity levels (Yamashita et al. 2007), we hypothesized that the microexon insertion could physiologically modulate DAAM1-mediated actin polymerization specifically in neurons. To test this hypothesis, we first purified the C-terminal part of the human DAAM1 protein with and without the microexon (Figure 2a). This region (FH2-COOH) encompasses the entire FH2 domain and has been previously shown to be functional in vitro (Lu et al. 2007). After selecting the optimal conditions for the experiment (Figure S1b-d), we compared the actin polymerization activity of both splice isoforms. Remarkably, the isoform without microexon (FH2-COOH (-MIC)) exhibited a significantly higher actin polymerization rate (Figure 2e,f).

**Figure 2.**
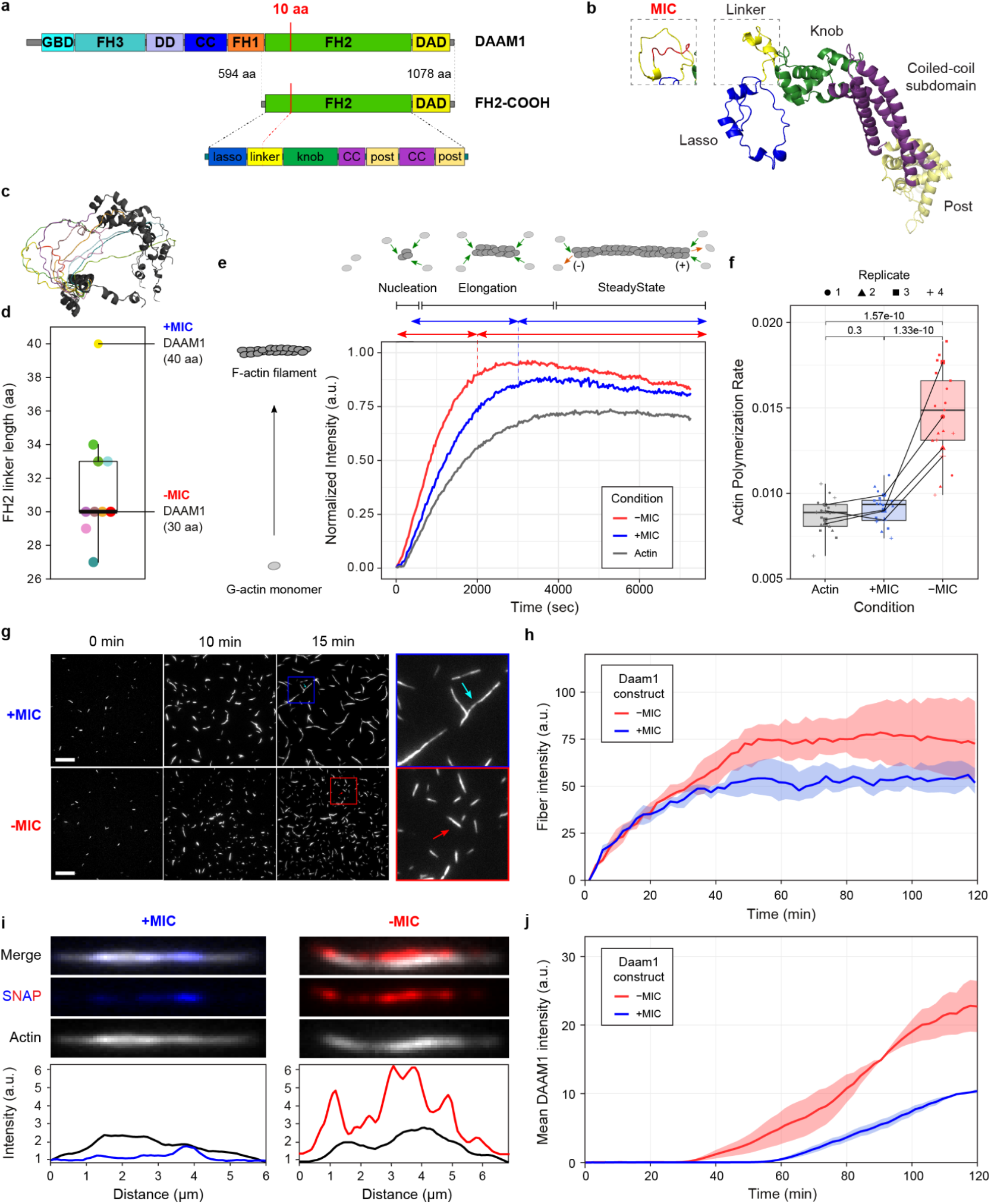
Splice variants of the DAAM1 FH2-COOH fragment differentially modulate actin dynamics. (a) Schematic representation of DAAM1 domains (top) and subdivisions within the FH2 domain (bottom). FH2-COOH corresponds to the purified protein fragment. Domain nomenclature as in Figure le. (b) Structure of the FH2 domain with (left) or without (right) microcxon inclusion. Subregions of the lasso, linker, coiled-coil as well as knob and post regions share the same color-code as in (a), (c) Structural comparison of the FH2 linker regions among all human formin proteins. Loop color corresponds to the formin color coding from Figure Sia. Structures were obtained from AlphaFold2 and visualized using the PyMoL program, (d) Lengths of the FH2 linker regions across formin proteins. The linker color corresponds to the structures in (c and Sia), (e) Pyrene actin polymerization assay. Each curve is the average of five technical replicates. Schematic representations of the nucleation, elongation, and steady-state phases of actin polymerization from the “Actin” profile are represented above, (f) Actin polymerization rates from four independent experiments. Rates correspond to the slope of the curves at 50% assembly (Doolittle, Rosen, and Padrick 2013). Data points correspond to technical replicates from four independent experiments. P-values from ANOVA with replicate and category as factors, (g) Representative images from time series of fluorescence micrographs showing F-actin fibers stained with SiR-Actin in the presence of microexon-containing (+MIC) or non-containing (-MIC) splice variant of the FH2-COOH fragment of human DA AM I. 2 pM Actin and 200nM DAAM1 fragments were used. Scale bar: 10 pm. On the right, zoom in on the regions of interest depicted in the 15-minute time point. Arrows indicate differences in actin fiber morphology, (h) Quantifying total fiber intensity through the experiments for 0.2 pM actin and 200 nM DAAM1 fragments. Thick lines represent the median from four independent experimental replicates, and the dispersion corresponds to the first and third quartiles. Statistical tests can be found in Figure S2i. (i) Dual-color T1RF microscopy of actin fibers obtained after 50 min using 0.2 pM actin and 200 nM SNAP-tagged proteins. The upper panel shows representative images of the actin fibers considered for the analysis. The lower panel displays the fluorescence intensity distribution along the fibers (black indicates actin, red and blue +MIC and -MIC proteins, respectively), (j) Quantification of protein fluorescence intensity along actin fibers as a function of time using 0.2 pM actin and 200 nM SNAP-tagged proteins. Thick lines represent the median from five independent experimental replicates, and the dispersion corresponds to the first and the third quartiles. Statistical tests for selected time points can be found in Figure S3f.

To further characterize the impact of Daam1-MIC in actin dynamics and structure, we performed Total Internal Reflection Fluorescence (TIRF) microscopy experiments to directly visualize the actin network, which revealed a much more complex picture of the functional differences between the two isoforms (Figures 2g and S2a). Consistent with the previous experiment, the isoform lacking the microexon exhibited an overall higher actin polymerization activity, as assessed by total intensity quantification (Figure S2b). Moreover, TIRF experiments showed that the difference between the two proteoforms was not solely in polymerization activity, but they also had a qualitatively different behavior, resulting in distinct organizations of the actin network (Figures 2g and S2a-c). Detailed comparison of the spatio-temporal evolution of the actin network’s morphology at the level of individual fibers (i.e., bundles of actin filaments) showed that the microexon-containing isoform led to an increased average length of the actin fibers (Figure S2e) and of their branches (Figure S2f), with a trend to increase in crossing junctions between fibers (Figure S2g) resulting in the formation of multiple branches (Figure S2h).

Previous studies also suggested that DAAM1 is a potent actin-bundling protein (Jaiswal et al. 2013). Remarkably, microexon removal resulted in significantly increased fiber fluorescence, indicating higher filament bundling activity for this isoform (Figures 2h and S2i). This suggests that the splice variants of DAAM1 interact with actin differently, in terms of kinetics and/or affinity, which could explain the behaviors described above. Thus, we next investigated whether the inclusion and skipping isoforms exhibited differences in binding to actin fibers. To do so, we performed dual-color TIRF microscopy experiments using SNAP-tagged DAAM1 FH2-COOH fragments (Figure S3a-c). Both SNAP-tagged protein isoforms showed strong co-localization with actin fibers (Figure 2i and S3d). However, analysis of individual actin bundles revealed a significantly higher amount of skipping isoform bound to actin fiber (Figures 2i,j, and S3e,f). This is likely due to the intrinsic activity of the isoform, as no major differences in dimerization ability were observed between the two isoforms based on the FPLC profiles (Figure S3a). In summary, these results demonstrate that microexon inclusion strongly impacts the functionality of the FH2 domain of DAAM1, both quantitatively and qualitatively. Microexon skipping increases actin polymerization rate and bundling, forming thicker fibers, while its inclusion promotes actin fiber elongation, creating more flexible and versatile actin networks and resulting in an overall more complex topology.

### Daam1-MIC deletion leads to increased neuronal excitability in glutamatergic neurons

To begin investigating the relevance of Daam1-MIC in neuronal differentiation and function, we deleted the microexon in mouse embryonic stem cells (ESCs) using the CRISPR-Cas9 system (see Methods). We selected three independent knockout (KO) and control ESC clones and differentiated them in vitro into glutamatergic neurons (Figure 3a) (Bibel et al. 2004; 2007). RT-PCR assays confirmed the precise microexon deletion in KO cells, with no associated mis-splicing during differentiation, whereas control cells displayed prominent microexon inclusion soon after the plating of neuronal precursor cells (DIV0) (Figures 3b and S4a).

**Figure 3.**
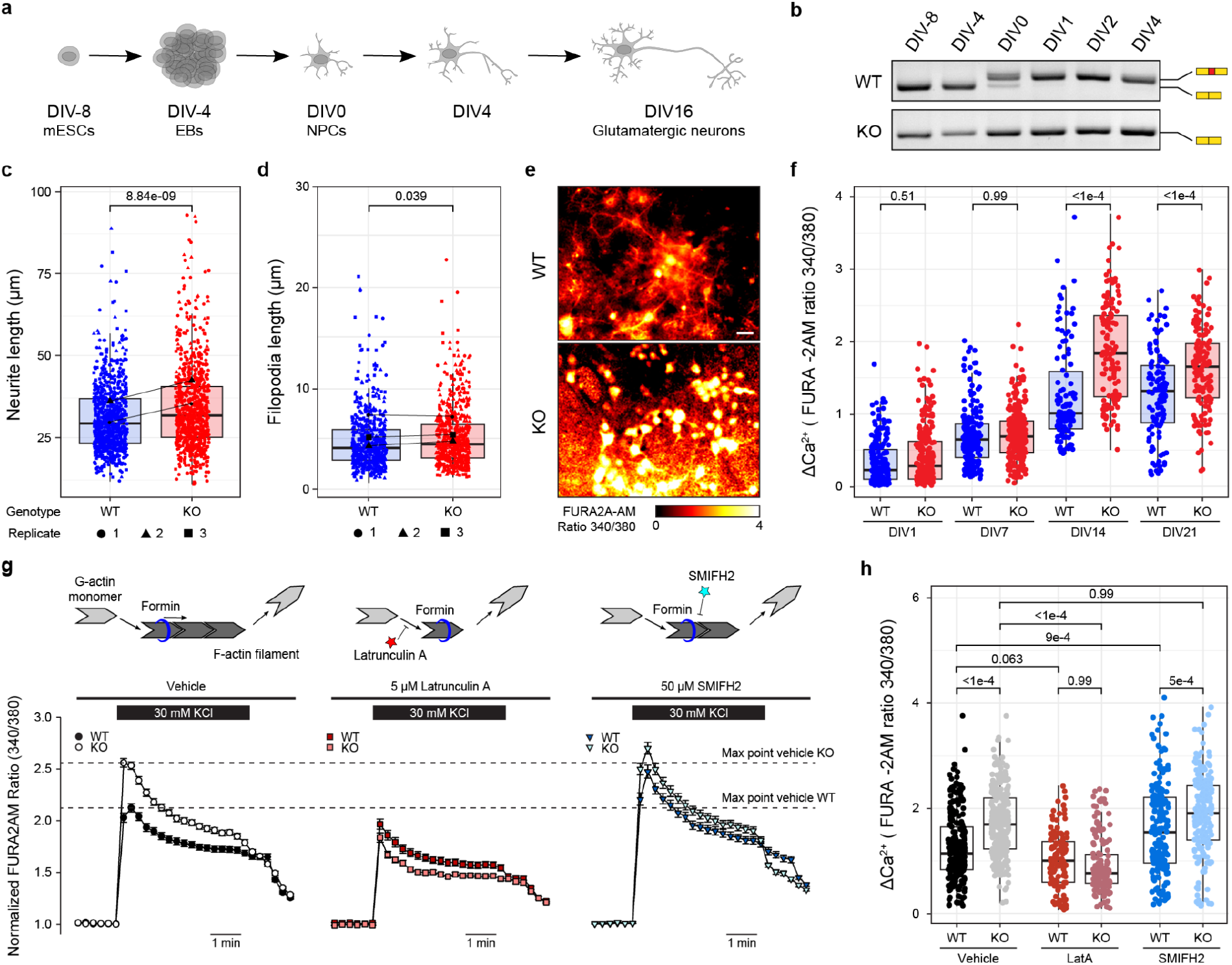
Daaml-MIC removal increases calcium flux in glutamatergic neurons. (a) Schematic representation of the neuronal differentiation protocol. DIV. day in vitro, (b) RT-PCR assays of Daaml-MIC inclusion during neuronal differentiation in representative WT and KO cell lines. (c,d) Distributions of the lengths of the longest neurites (c) and individual filopodia (d) of neuronal precursors (DIV0+4h). P-values from two-way ANOVA tests with replicate and genotype as factors, (e) Representative images of the calcium imaging experiment performed using FURA-2AM in mature neuronal cells (DIV21) depolarised with 30 mM KC1 isotonic solution. Scale bar: 10 pm. (f) Distributions of calcium influx in WT and KO cell lines at various differentiation time points, based on FURA-2AM ratio. Neurons were stimulated with a 30 mM KC1 isotonic solution. P-values from ANOVA Tukey’s test, (g) Effects of the actin polymerization inhibitor Latrunculin A (LatA) and the small molecular inhibitor of formin FH2 domains (SMIFH2) on calcium currents in differentiated glutamatergic neurons at DIV14-21, Data corresponds to three independent experiments. Top: schematic representation of the mode of action of the actin-modulating drugs, (h) Distributions of calcium influx in WT and KO cell lines upon different treatments as depicted in (g), based on FURA-2AM ratio. P-values from Kruskal-Wallis test.

Since DAAM1-driven actin polymerization is important for neurite morphology and filopodia integrity (Jaiswal et al. 2013; Matusek et al. 2008; Szikora et al. 2017), we first assessed the impact of the microexon removal at early stages of neuronal differentiation (Figure S4b). Daam1-MIC removal led to a significant increase in neurite length (Figure 3c), consistent with the increased actin polymerization observed in vitro (Figure 2). Additionally, we observed a slight increase in filopodia length in KO compared to WT cells (Figure 3d), but no significant changes in filopodia number (Figure S4c). Immunostaining of DAAM1, actin, and α-tubulin showed no major differences in the protein signal within the growth cones between the two genotypes (Figure S4d,e), with only a small relative reduction of DAAM1 and actin in more distal filopodia in KO cells (Figure S4f,g).

Despite these effects on neuritogenesis, microexon removal did not impair the ability of neurons to differentiate, as both WT and KO cell lines successfully generated neuronal networks and formed functional synaptic connections. However, mature KO neurons (DIV14 and DIV21) exhibited a significant increase in Ca2+ influx specifically upon depolarization (Figure 3e,f). To further investigate this phenotype, we performed chemical treatments with modulators of various aspects of actin dynamics (Figure 3g). Reduction of F-actin polymerization accomplished by sequestering available G-actin monomers with Latrunculin A (LatA) resulted in a strong reduction of Ca2+ influx in both WT and KO cell lines, abolishing the differences between the two genotypes (Figure 3g,h), and suggesting that these differences largely depend on actin polymerization dynamics. Next, we applied a small molecular inhibitor of formin FH2 domains (SMIFH2) that selectively inhibits formin-driven actin polymerization without impairing polymerization by other actin-binding proteins. Exposure of neuronal cultures to SMIFH2 increased Ca2+ influx in WT cell lines, mimicking the KO phenotype (Figure 3g,h), and suggesting that microexon removal in neurons leads to inappropriate activity of the FH2 domain of DAAM1. Finally, the increase in Ca2+ influx observed in Daam1-MIC KO neurons was neither due to differences in basal calcium levels (Figure S4h), to DAAM1 isoform localization to the synapses (Figure S4i) nor to changes in protein expression (Figure S4j). Altogether, these results suggest that Daam1-MIC removal in mature neurons leads to increased Ca2+ influx upon depolarization due to effects in actin polymerization dynamics.

### Daam1-MIC KO adult mice exhibit learning defects

Next, we generated a Daam1-MIC KO mouse line by ESC blastocyst microinjection (Figure S5a and see Methods). In contrast to the full gene KO (Nakaya et al. 2020), Daam1-MIC KO was not lethal, and mutant mice were born with expected Mendelian ratios (Figure S5b), had normal weight (Figure S5c) and did not display gross morphological abnormalities. To assess putative effects during the pre-weaning period, we first performed a neurodevelopmental behavioral screen (Figure S5d) (Feather-Schussler and Ferguson 2016; Roper et al. 2020). These experiments showed no major differences between KO and WT siblings in basic reflexes (Figure S5e-g), balance and motor coordination (Figure S5h-k). Mild but significant increases were observed in KO mice in forelimb and hindlimb strength (Figure S6a,b), and motor development-dependent walking (Figure S6c). Interestingly, we also observed a decrease in the performance of the homing task (Figure S6d), indicating delayed psychomotor development, but we found no significant gross morphological differences in dentate gyrus size (Figure S6e,f).

We then assessed a range of potential defects in adult mice. Remarkably, whereas no major differences were observed in motor function and balance (Figure S7a-f) or in anxiety-related phenotypes (Figure S7g-k), we found that learning and memory formation capabilities were robustly affected upon Daam1-MIC removal. First, KO mice of both sexes showed significantly impaired recognition memory, evaluated using the novel object recognition (NOR) task (Figure 4a,b), which depends on the prefrontal cortex and hippocampal circuit responsible for cognitive processing (Leger et al. 2013; Warburton and Brown 2015; Lueptow 2017). In contrast, no significant differences in total exploration time were observed during the familiarization or test phases of the experiment (Figure S8a-b). Second, we assessed spatial memory by employing a modification of the Morris Water Maze (MWM) task (Figures 4c,d and see Methods) (Vorhees and Williams 2006). This test showed impaired spatial learning specifically in KO males between days 4 and 8 (Figures 4e,f and S8e,f). This impairment was not due to motor or motivation defects, as no differences were detected during the cued session when the platform was visible (Figure 4e,f). At the molecular level, no differences between genotypes were found in the expression of cFOS or EGR1 protein levels (Figure S8g-j) or DAAM1 protein expression levels in mouse cerebellum, motor cortex, or other tissues (Figure S8k-m). Altogether, these results suggest that Daam1-MIC removal negatively impacts memory formation in adult mice, a defect that cannot be attributed to impaired motor function or anxiety-related behaviors.

**Figure 4.**
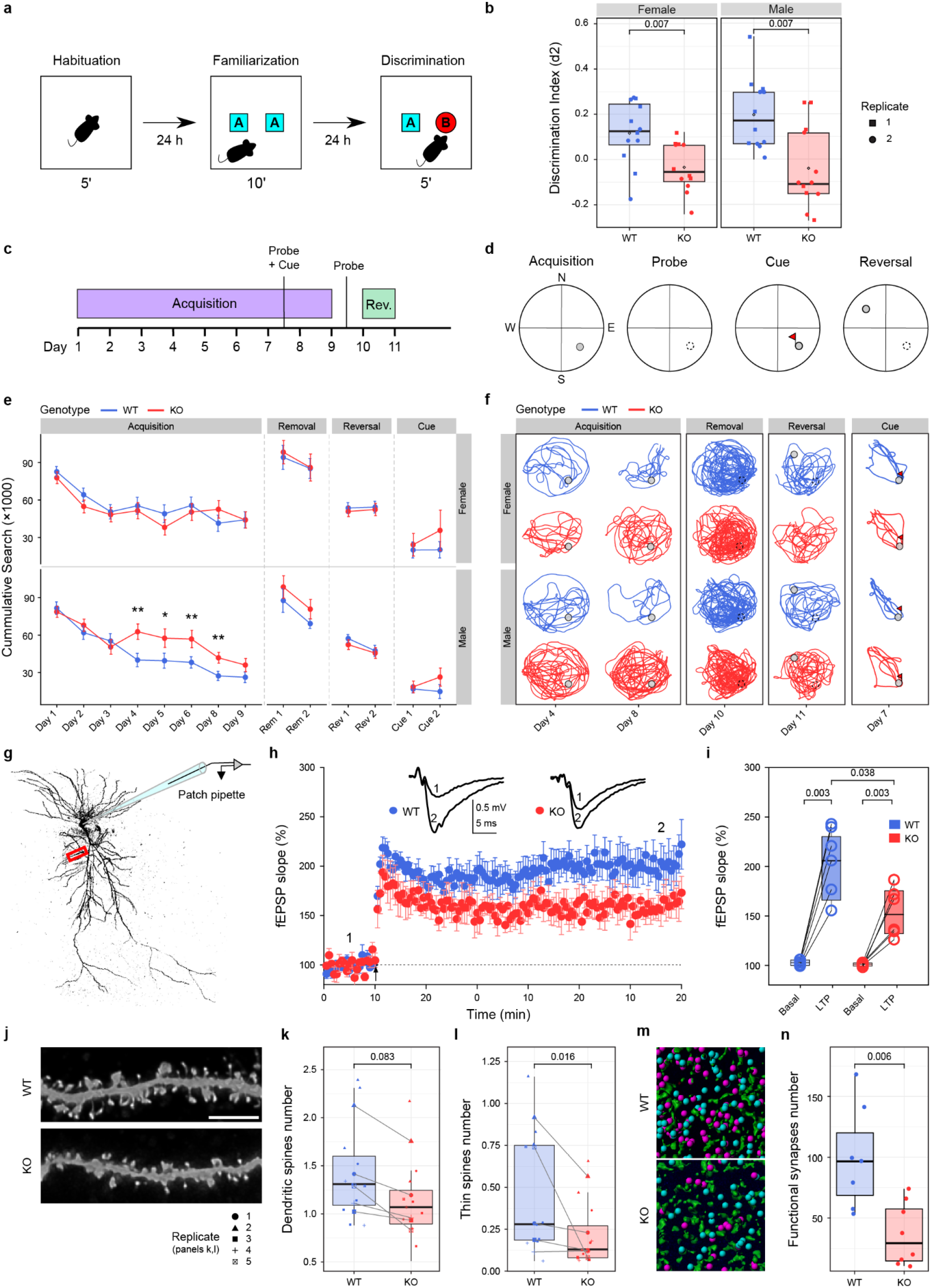
Daaml-.MIC removal causes learning impairments by modulating the hippocampal LTP. (a) Schematic representation of the Novel Object Recognition (NOR) experiment. The experiment consists of three sessions: habituation, familiarization, and discrimination (lengths of 5 min, 10 min, and 5 min, respectively). Each session was separated by 24 h periods, (b) Distribution of discrimination index (d2) values quantified during the NOR discrimination phase. The discrimination index corresponds to: (Novel Object Exploration Time - Familiar Object Exploration Time)/ Total Exploration Time. Each data point describes the performance of one animal. P-values from Wilcoxon rank-sum tests. (c,d) Schematic representation of the Morris Water Maze (MWM) protocol (c) and platform set-ups (d) (details in Supplementary Methods), (e) Cumulative search index during the acquisition, removal, reversal, and cued phases for females (top) and males (bottom) based on (Tomas Pereira and Bunveil 2015). Distributions of values for days 4 and 8 are shown in Figure S7f. P-values from two-way ANOVA tests with replicate and genotype as factors, (f) Representative trajectories of performances of all six animals from one trial of experimental replicate 1 during each phase of the MWM. (g) Representative maximum-intensity confocal image of a dye-filled neuron after deconvolution. The red box marks the dendrite region analyzed, (h) Field excitatory postsynaptic potentials (fEPSPs) were recorded in the CAI dendritic layer in response to CAI Schaffer collateral stimulation. LTP was induced by theta burst stimulation (TBS; 10 bursts of 5 pulses at 100 Hz, with an interval of 200 ms between bursts). LTP temporal plot of fEPSP values in KO (red circles, n= 6, N= 5) compared to WT mice (blue circles, n = 5, N = 3) after TBS at 60 min. Insets of traces in the plots represent average fEPSPs recorded during periods indicated by corresponding numbers in the graph (1 and 2). (i) Boxplot of LTP after TBS at 60 min. P-values from Welch’s t-tests. 0) 3D reconstructions of pre-(Synapsin 1/2, green) and post-(PSD95. blue and pink) synaptic markers in PD22 CAI hippocampal sections. Pink spots represent postsynapses at <0.5pm from the presynapses (functional synapses) and blue ones at >0.5pm. (k) Distributions of the percentage of functional synapses normalized to the control mean for WT and KO animals. P-values from Wilcoxon rank-sum tests. (I) Representative maximum-intensity confocal images of dye-filled dendrites and dendritic spines after deconvolution. Scale bars: 5 pm. (m.n) Number of total dendritic spines (m) and thin spines (n). N = 5 WT and 6 KO mice, 100 pm of dendrites per cell. One dot corresponds to one neuron analyzed. Lines represent the relation between the averaged values from the animals analyzed in experimental replicates. P-valucs from Wilcoxon rank-sum tests.

### Daam1-MIC removal impacts LTP and leads to reduced synaptic spines

To obtain further mechanistic insights into these phenotypes, we next investigated how Daam1-MIC deletion affected synaptic function and plasticity in hippocampal slices from adult males. Whole-cell voltage-clamp recording in CA1 pyramidal cells of the hippocampus showed a trend for a higher number of spikes in response to suprathreshold depolarizing current (Figure S9a), suggesting a mild increase in the firing rate compared to WT neurons (Figure S9b). We observed no changes in the set of parameters that describe intrinsic membrane properties (rheobase, membrane resistance, membrane potential, and threshold action potential; Figure S9c-h), indicating that ionic permeability was not affected, and there were no changes in membrane conductance in KO neurons.

We next studied the effect of Daam1-MIC removal on glutamatergic CA3-CA1 synapses. We observed a strong and significant decrease in field Excitatory PostSynaptic Potential (fEPSP) based on induced Long-Term Potentiation (LTP) using a Theta Burst Stimulation (TBS) in KO neurons (Figure 4g-i). To explore possible mechanisms behind this decrease, we then examined basal synaptic transmission in the CA1 region (input-output relationship of excitatory postsynaptic currents (EPSC)) and the presynaptic release of neurotransmitters, by applying Paired-Pulse Facilitation protocol (PPF) and recording miniature EPSCs (mEPSCs). These analyses did not show differences between WT and KO mice (Figures S9i-p), suggesting no changes in basal excitatory synaptic transmission at hippocampal (CA3-CA1 neuron) synapses, and ruling out major presynaptic alterations associated with the LTP deficit in KO slices. In contrast, subcellular analysis of pre- (Synapsin 1/2) and post- (PSD95) synaptic markers in the CA1 region of the hippocampus at an earlier developmental time point (P21) showed a striking decrease in the number of PSD95-positive puncta, but no differences in Synapse 1/2 puncta (Figure 4j,k, and S9r,s). Interestingly, biocytin labeling of neurons in adult mice revealed a significantly reduced number of thin immature dendritic spines (Figure 4l-n), but not of stubby and mushroom-like spines (Figures S9t). Therefore, altogether, these results indicate a severe impairment of the postsynaptic densities likely explaining the decrease in LTP in Daam1-MIC KO mice.

### Daam1-MIC controls the RHOA/ROCK signaling pathway activity

Taken together, these results suggest that Daam1-MIC deletion may alter molecular pathways associated with actin cytoskeleton that are essential for the proper formation of dendritic spines and LTP. As mentioned above, the RHOA/ROCK signaling cascade is essential for controlling the actin cytoskeleton-driven dendritic spine formation and LTP (Zhang et al. 2021), and DAAM1 has been shown to directly interact with RHOA (Habas, Kato, and He 2001; Higashi et al. 2008; Kühn and Geyer 2014). Therefore, we next investigated whether microexon inclusion/skipping changes this protein-protein interaction. As previously described (Liu et al. 2008), the removal of the DAAM1 autoinhibitory domain (DAD) (Figure S10a,b) increased binding with RHO proteins, augmenting Co-IP efficiency (Figure S10c,d). Using this construct, Co-IP assays showed a significant ∼40% decrease in the interaction between DAAM1 and RHOA upon microexon removal (Figures 5a-b and S10d-f). Remarkably, a significant increase of similar magnitude in RHOA activity in protrusions of mature KO neurons (Figures 5c,d and S10g) was observed by acceptor photobleaching of a RhoA FRET-based biosensor (Fritz et al. 2013).

**Figure 5.**
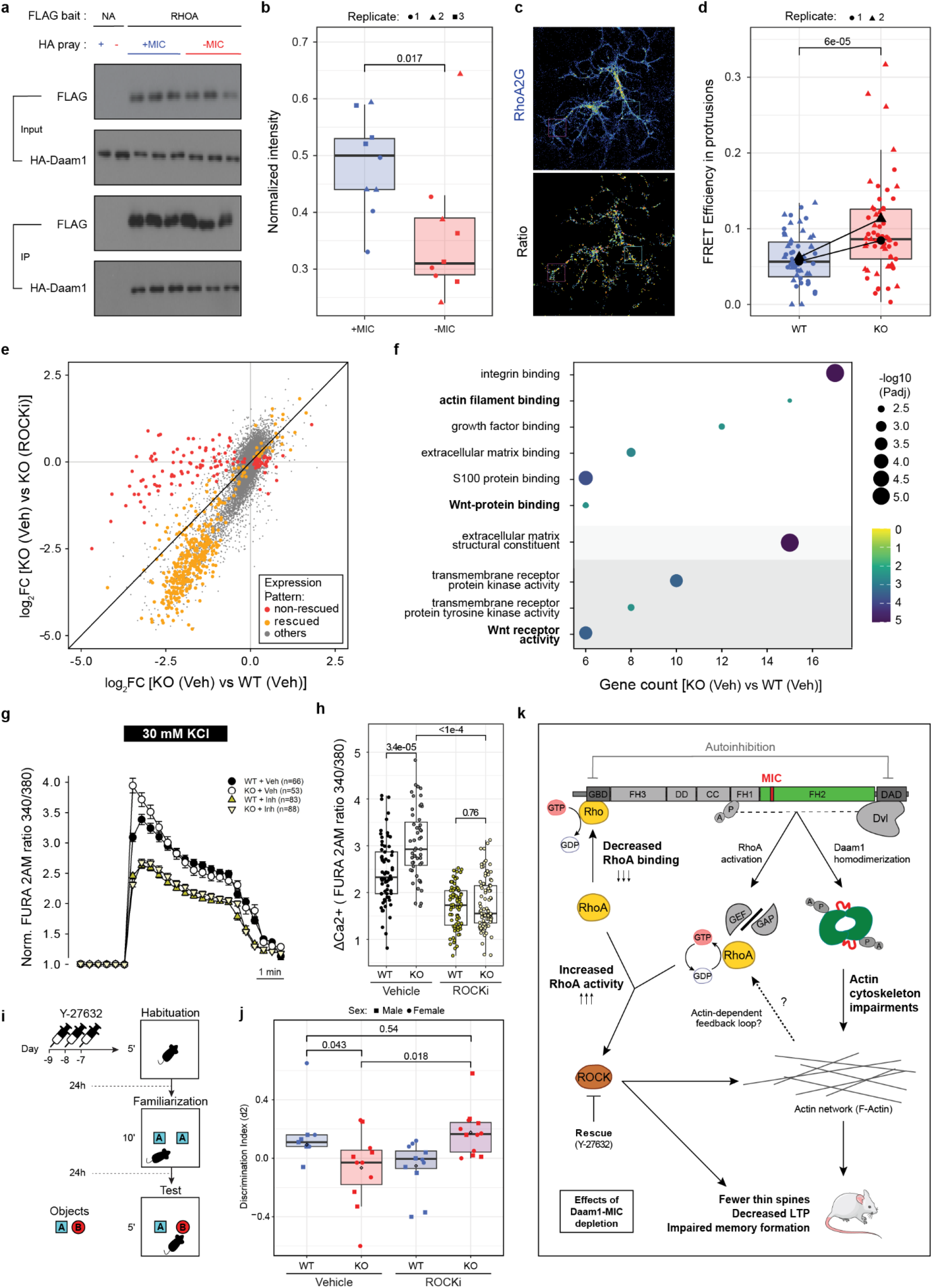
Loss of Daaml-MIC impairs RHOA/ROCK signaling cascade. (a) Co-immunoprecipitation of DAAM1 with its interactor RHOA. 293T cells were transfected with HA-tagged DAAM1-ADAD expression constructs, with (+MIC) or without (-MIC) the microexon, together with 3xFlag-tagged RHOA expression vector. Immunoprecipitation was performed with anti-FLAG antibody, (b) Quantification of the interaction between DAAM1-ADAD constructs with (+MIC) or without microexon (-MIC) and RHOA in the Co-IP experiment presented in panel (a) and SlOe.f. P-values from Wilcoxon rank-sum tests, (c) Confocal micrographs representing RhoA2G distribution (top) and RhoA activity (sensor ratio: bottom) in mature WT neuronal cells (DIV21). Boxes represent examples of regions of interest. Scale bar: 50 pm. (d) Comparison of the RhoA2G biosensor FRET efficiency between WT and KO cell lines using fluorescent microscopy. One dot represents FRET efficiency in one neuron. P-values from two-way ANOVA tests with replicate and genotype as factors, (e) Scatter plot showing a comparison of changes in gene expression levels (Iog2 fold changes) between WT and KO cells in the control treatment (Veh) (X-axis) vs. changes between KO control cells and KO cells treated with ROCK inhibitor (Y-27632) (Y-axis). Each data point corresponds to a gene, which is colored according to their “rescue” group: orange, misregulated expression in the KO rescued by the ROCKi; red, misregulated expression in the KO not rescued by the ROCKi; grey, other genes (see Methods), (f) Molecular Function GO term enrichment for genes with a rescue pattern upon ROCK inhibitor application (orange in (e)). (g,h) Distributions of calcium influx measurements in WT and KO cell lines upon ROCK inhibitor treatment based on FURA-2AM ratio. P-values from Kruskal-Wallis test, (i) Schematic representation of the Novel Object Recognition (NOR) experiment performed to test the effect of ROCK inhibition. Three consecutive injections of ROCK inhibitor or vehicle were applied one week before the NOR at days -9, -8, and -7. (j) Distribution of discrimination index values in the different conditions and for each genotype. P-values from Wilcoxon rank-sum tests, (k) Summary of the observed phenotypes upon Daam I -MIC removal and their relation to the RHOA/ROCK signaling cascade (based on (Habas, Kato, and He 2001; Liu et al. 2008; Schonichcn and Geyer 2010)). DVL binds to DAAM1 releasing its autoinhibition, which is further facilitated by interaction with RHOA. Active DAAM1 leads to RHOA activation through an unknown mechanism involving DAAMl’s C-terminal part (dashed lined) and is not dependent on direct binding RHOA-DAAMI interaction. Microexon removal decreases RHOA binding to DAAM 1 ‘s N-terminal region. We propose that this leads to lower hydrolysis of RhoA-GTP, increasing the pool of active RHOA and in turn hyperactivating the RHOA/ROCK signaling cascade. Arrows indicate the directionality of the event. The dotted arrow indicates a potential, unknown feedback loop from actin polymerization to ROCK pathway activation. GAP - GTPase-activating protein, GEF - guanine nucleotide exchange factor, GTP - Guanosine triphosphate, GDP - Guanosine diphosphate.

Based on these observations, we hypothesized that increased RHOA activity upon Daam1-MIC removal leads to hyperactivation of the RHOA/ROCK signaling cascade, which, in turn, causes some of the observed phenotypes. To test this hypothesis, we used an inhibitor of ROCK (Y-27632), a direct downstream effector of RHOA. First, application of Y-27632 to differentiated neurons rescued the expression alterations of most differentially expressed genes between the KO and WT cells (Figures 5e and S11a-c). This included multiple genes related to the function of the RHOA/ROCK signaling cascade, the Wnt-pathway, and the regulation of the actin cytoskeleton (Figures 5f and S12). Second, applying Y-27632 reverted the difference in Ca2+ influx between WT and KO neurons (Figure 5g,h). Lastly, injection of Y-27632 intraperitoneally in mice significantly increased the discrimination index of KO animals of both sexes, suggesting at least a partial rescue of the memory impairments (Figures 5i,j and S11d-i). Moreover, ROCK inhibitor treatment caused a decrease in the discrimination index in WT animals, further supporting genotype-specific differences in response to changes in RHOA/ROCK signaling (Figure 5j). Altogether, these results suggest that Daam1-MIC is important for the correct molecular interplay that controls the RHOA/ROCK signaling cascade regulation (Figure 5k), which is involved in precisely controlling memory formation.

## Discussion

Tight regulation of actin cytoskeleton polymerization and its associated molecular pathways by actin-binding proteins is crucial for multiple neurobiological processes. Here, we found that Daam1, the formin gene with the highest expression in mature neurons, harbors a microexon that is highly neural-specific and evolutionarily conserved across vertebrates. Interestingly, we show that correct regulation of Daam1-MIC is crucial at multiple levels. Firstly, microexon inclusion structurally extends the length of the linker region of the FH2 domain, modulating its actin polymerization capabilities, quantitatively and qualitatively, in controlled in vitro assays. Secondly, Daam1-MIC removal led to a decrease in DAAM1-RHOA interaction and enhanced activity of the RHOA/ROCK signaling pathway in neurons, which is known to influence multiple cellular processes, including actin cytoskeleton dynamics. Lastly, deletion of the microexon led to diverse phenotypes in mice, including a decreased number of dendritic spines, reduced LTP, and defects in memory formation.

The inclusion of Daam1-MIC in neurons makes the length of the DAAM1’s FH2 linker region an outlier with respect to all other formins.

Given that the linker region serves as a hinge in FH2 dimers (Schönichen and Geyer 2010), its uniquely large length likely results in a much more flexible dimer, which we show to bind less to actin as compared to the non-neural form lacking the peptide encoded by the microexon. Previous studies (Yamashita et al. 2007; Moseley, Maiti, and Goode 2006) reported that the long linker of DAAM1 was associated with this protein’s unusually low actin polymerization rate and that its shortening enhanced polymerization. We recapitulate these findings and further connect them with a splicing-driven and physiologically occurring event that modulates DAAM1’s function in a tissue-dependent manner. Moreover, our TIRF results revealed a much more complex scenario in which microexon inclusion does not simply lead to a reduced polymerization rate but to a dramatic change in how DAAM1 interacts with and remodels the actin cytoskeleton. In particular, we observed that microexon inclusion significantly decreased the actin-bundling capabilities of DAAM1, probably as a direct result of a lower affinity to actin. Moreover, a detailed structural characterization of actin bundles showed that the microexon-containing isoform produced consistently longer and more branched fibers, likely due to its less constrained binding to actin. Although these results are based on in vitro reconstituted assays, it is tempting to speculate that, *in vivo*, the inclusion or exclusion of the microexon may similarly modulate DAAM1’s behavior with actin and hence its function. In line with this idea, at the early stages of neuronal differentiation, we observed a significant increase in neurite length in Daam1-MIC KO neurons, consistent with enhanced actin polymerization.

In our mouse model, Daam1-MIC removal resulted in memory impairments, plausibly due to decreased LTP caused by a reduced number of dendritic spines. In particular, we observed a decreased number of thin, immature spines characterized by their dynamic structure, and considered “learning spines” as they can be consolidated into more stable mushroom spines ((Lamprecht 2021; McLeod and Salinas 2018)). What might be the molecular bases of these cellular phenotypes? Rho-family GTPases, including RHOA, are key players in memory formation by regulating spine morphology, at least partially via their role in regulating actin cytoskeleton (Zhang et al. 2021). Evidence suggests that RHOA activity modulates immature spine pruning (Bolognin et al. 2014; Zhang et al. 2021). For instance, expression of constitutively active RHOA in hippocampal neuronal cultures or brain slices resulted in simplified dendritic trees and reduced spine densities (Nakayama, Harms, and Luo 2000; Impey et al. 2010). Conversely, RHOA inhibition specifically increases the number of immature spines (Nakayama, Harms, and Luo 2000). Additionally, to further establish the link between Daam1-MIC deletion, RHOA/ROCK signaling and some of the observed phenotypes, we used Y-27632, a specific ROCK inhibitor. Y-27632 treatment of differentiated glutamatergic neurons decreased calcium influx upon depolarisation abolishing the observed differences between the genotypes. Moreover, administration of the inhibitor recovered most of the gene expression alterations observed upon microexon removal and was sufficient to revert the memory deficiencies observed in the NOR task performed by the Daam1-MIC KO mice. Altogether, these results suggest a causative upregulation of the RHOA/ROCK signaling cascade upon microexon removal.

How could Daam1-MIC removal lead to upregulation of the RHOA/ROCK signaling cascade? DAAM1 is known to directly interact with RHOA, with a binding preference towards active GTP-RHOA (Liu et al. 2008; Habas, Kato, and He 2001). Interestingly, even though the microexon does not lie in the proximity of the RHOA interacting domain (N-terminal GDB domain), our Co-IP assays showed a significant decrease in the interaction between DAAM1 and RHOA upon microexon removal. Could this lead to the observed increase in overall RHOA activity? The exact mechanism of DAAM1-dependent activation of RHOA is unknown, but it does not rely on the direct interaction of DAAM1 with RHOA and it is instead believed to occur through the recruitment of the GEF-RHOA complex to DAAM1 (Habas, Kato, and He 2001). On the other hand, in Diaphanous-related formins, such as DAAM1, the direct binding of RHOA-GTP to the GDB domain is followed by hydrolysis to RHOA-GDP, contributing to formin activation (Kühn and Geyer 2014; Higashi et al. 2010). Since DAAM1’s activation is highly dependent on DVL proteins (Liu et al. 2008), it is possible that reduced interaction of RHO-GTP with DAAM1’s GDB domain in Daam1-MIC KO cells mainly results in less RHOA deactivation and thus an increased pool of active RHOA-GTP available (Figure 5k). Another unexplored possibility is that alterations in actin dynamics upon Daam1-MIC depletion indirectly affect RHOA activity through an unknown mechanism (Figure 5k), independently or in addition to the change in DAAM1-RHOA protein interaction. Given the importance of RHOA/ROCK signaling in modulating actin dynamics, it is plausible that such feedback loops exist.

In summary, through a comparative analysis of formin proteins, we found that Daam1 is subject to tight post-transcriptional regulation by a highly neural-specific and evolutionarily conserved microexon. Daam1-MIC inclusion directly impacts actin polymerization via modulation of the FH2 domain activity. Moreover, through an in-depth multi-level characterization of this microexon, we unveiled DAAM1 as a key new player in regulating neurotransmission and memory formation.

## Materials and Methods

### Computational analysis formins and characterization of DAAM1 and RT-PCR assays

To investigate the expression of formins across tissues and during neuronal differentiation (Fig. 1a), we used mouse (mm10) data from VastDB (Tapial et al. 2017). In addition, we extracted the mouse and/or human neural-regulated exons among the 15 mammalian formin genes from Table S2 in (Irimia et al. 2014) and used VastDB to extract the exon’s protein location and inclusion levels for the six identified neural regulated exons shown in Fig 1b and 1c, respectively. The study of the evolutionary conservation of these six exons was carried out using the VastDB ortholog information and manual exon alignments among the human (hg38), mouse (mm10), cow (bosTau9), opossum (monDom5), chicken (galGal4), frog (xenTro9), zebrafish (danRer10) and shark (calMil1) genomes, downloaded from Ensembl. To investigate the regulation of Daam1-MIC by SRRM3/4 (Fig. 1i), we used data from human HEK293 cells overexpressing human SRRM4 (GEO: GSE112600 (Torres-Méndez et al. 2019)), mouse N2A cells upon Srrm3/4 knockdown (SRA: PRJNA47491 (Gonatopoulos-Pournatzis et al. 2018)), and zebrafish retinae extracted from Srrm3 KO 5 days post fertilization larvae (GEO: GSE180781 (Ciampi et al. 2022)). Crystal Structure of human DAAM1 FH2 domain (2z6e.pdb) was used to model the mice Daam1 FH2 domain with and without microexon using Robetta (http://new.robetta.org/) (Fig. 2b). Visualization was performed using the PyMoL program.

To validate the inclusion of Daam1-MIC through RT-PCR assays in mouse and zebrafish (Fig. 1h and throughout the manuscript), total RNA was extracted using the RNeasy Plus Mini kit (Qiagen, 74136), treated with DNase using TURBO DNA-freeTM Kit (Thermo Fisher Scientific), and reverse-transcribed with SuperScript III Reverse transcriptase (Invitrogen, 18080044) and oligo(dT)20 as advised by the manufacturer. GoTaq G2 Flexi DNA Polymerase enzyme (Promega, M7806) was used to amplify Daam1-MIC’s flanking exons and PCR assays were performed with 25 cycles, 54°C annealing temperature and 120 sec extension. The PCR product was size-fractionated using 2.5% ultrapure agarose (Invitrogen, 16500500) in SB buffer (36.4 mM of Boric acid and 10 mM NaOH), and detected using SYBR Safe (Life Technologies, S33102). Primer sequences used in this study are provided in Table S1.

### Cloning of Daam1 FH2 domain, protein expression and purification

To clone the functional FH2 domain of DAAM1 (residues 594-1077), we have used RNA extracted from Human Embryonic Kidney 293 (HEK293T) and SH-SY5Y cells. RNA samples were extracted from 1 million cells using the RNeasy Mini Kit (Qiagen, 74136) and treated with TURBO DNA-freeTM Kit (Thermo Fisher Scientific, AM2238). Afterwards, 1 μg of RNA was reverse transcribed into cDNA using High-Capacity cDNA Reverse Transcription Kit (Thermo Fisher Scientific, 4368814). Primers used to amplify the FH2 domains of DAAM1 are provided in Table S1. The PCR product was cloned in pEMT33 (for actin polymerization assay and TIRF microscopy experiments) and in C145 pCoofy17 (for dual-color TIRF microscopy experiments), both vectors have been kindly provided by the Sebastian Maurer lab (CRG).

These constructs were expressed as fusion proteins with N-terminal IgG tag plus C-terminal Strep-tag in E. coli strain BL21-CodonPlus(DE3)-RIL strain (Agilent Technologies, 230245). Cells were cultured at 37°C until OD600nm reached 0.5, after which the media was cooled to 18°C and Isopropyl β-d-1-thiogalactopyranoside (IPTG; ROTH, 2316.1) was added to a final concentration of 0.1 mM to induce protein expression. Bacterial cultures were grown overnight at 18°C and subsequently harvested by centrifugation at 4°C (15 min, 4000 rpm) and disrupted by sonication. Sonication was conducted in HEKG 10 buffer (20 mM HEPES, pH 7.4, 1 mM EDTA, 50 mM KCl, 10% glycerol) with 1 mM DTT supplemented with cOmplete EDTA-free Protease Inhibitor Tablet (Sigma, 11873580001). After centrifugation (Eppendorf 5810R Centrifuge) at 4°C (20 min, 20000 rpm), proteins were purified using StrepTrap HP column (GE Healthcare, GE17-5248-01) at 4°C. The N-terminal tag was cleaved with His-C3 protease and the C-terminal tag with His-TEV protease (in house purified). Both proteases were further removed using Ni-NTA Agarose beads. Individual pooled DAAM1-FH2 proteins, with and without the microexon, were further purified by a passage through a Superose 6 column (GE Healthcare). Their concentration was evaluated using NanoDrop spectrophotometer (ThermoFisher) and their purity/integrity was assessed on SDS-PAGE, where coomassie staining (ThermoFisher, LC6065) was used for visualization.

### Pyrene actin polymerization assays and TIRF microscopy

Pyrene-labeled actin protein (Universal Biologicals) was prepared as suggested by the manufacturer. In short, actin was equilibrated for 1 h at 4°C in a G-buffer (5 mM Tris-HCl pH 8.0, 0.2 mM CaCl2) and spun for 30 min at 14 000 rpm before adding 10X Polymerization buffer (500 mM KCl, 20 mM MgCl2, 10 mM ATP) to a final 1X concentration. Actin assembly was measured in 60 μl reactions using 96 Well Black Polystyrene Microplates (Corning®, CLS3904). Pyrene fluorescence was monitored over time at 24°C at an excitation of 365 nm and emission of 407 nm in a fluorescence spectrophotometer (Tecan Infinite 200 PRO). Rates of actin assembly were calculated from the slope of the assembly curves at 50% polymerization (i.e., the time at which 50% of the actin has polymerized, or half time). As described by (Doolittle, Rosen, and Padrick 2013), we computed the half-time of each experiment based on the intensity values included in a specific interval. First, we calculated the average minimum intensity (average intensity of the 10 points closer to the minimum intensity) and the average maximum intensity (average intensity of the 10 points closer to tmax, where tmax is the average of the 10 points with the highest intensity). Then, we used these values to compute the lower and higher bounds of the intensity interval used for the half-time computation. The lower bound was defined as (0.5 - delta/2)*(avg_max_intensity - avg_min_intensity) + avg_min_intensity, while the higher bound was defined as (0.5 + delta/2)*(avg_max_intensity - avg_min_intensity) + avg_min_intensity (with delta=0.1, as recommended in (Doolittle, Rosen, and Padrick 2013)). Next, we used the points in each experiment with intensity values included between the lower and higher bounds to fit a linear model (intensity vs. time) and extracted the slope value returned by the fit. Finally, we derived the actin polymerization rate at half time by multiplying the slope value by a scaling factor (SF). SF was computed as (actin_conc - critical_conc)/(avg_max_intensity - avg_min_intensity), with actin_conc = x μM (actin original concentration as specified), critical_conc = k_minus / k_plus. k_minus and k_plus represented the on-rate and off-rate constants for filament assembly, and were set at 1.4 S–1 and 11.6 μM-1s-1, respectively, as suggested by (Doolittle, Rosen, and Padrick 2013)).

TIRF-based F-actin polymerization experiments were performed using manually assembled flow cells consisting of piranha-cleaned, silanized, PEG-passivated glass coverslips (22 X 22 mm #1.5 (i.e. 170 +/- 5 μm) from MARIENFELD (Cat # 0107052)) and a PLL-PEG passivated slide (Consolati et al. 2022). Flow cells were primed by flushing 55 μl of a solution of 5% Pluronic F-127 (Sigma-Aldrich, P2443) and then incubated for 10 min at room temperature. Next, flow cells were washed twice with 55 μl of kappa-Casein (1:100 in 1X G-buffer of 5 mg/ml stock in 1X Brb80 buffer). Absorptive paper (Whatman filter paper) was used to flush the chamber by holding the paper on the outlet side of the channel while adding the solution dropwise in the inlet side. Chambers were then equilibrated with G-buffer and the reaction mixture was loaded. The actin polymerization reaction mix contained the protein variant of interest (DAAM1-FH2 with or without microexon) and actin, which were diluted in freshly prepared buffer containing 5 mM Tris-HCl (pH 8.0), 0.2 mM CaCl2, 50 mM KCl, 2mM MgCl2, 2mM ATP, 2mM DTT, 1% (w/vol) glucose, 0.2 mM Brij-35, oxygen scavengers (180 mg/ml catalase (Merck, C40) and 752 mg/ml glucose oxidase (Serva, 22778.01)), 0.15% (w/vol) methylcellulose (Sigma-Aldrich, 4000 cP). Actin labeling was achieved using SiR-Actin (TeuBio, SC001). After loading the reaction mixture, flow cells were finally sealed with vacuum grease and placed onto the microscope stage for observation. To minimize the experimental variations between the different conditions to be compared (actin alone, actin with +MIC, and -MIC protein variants), experiments have been run in parallel, i.e. the three conditions were run simultaneously during each experiment. To this end, flow cells were divided into three separate channels using double-sided tape.

TIRF imaging was performed with a custom-assembled system (Cairn Research, Faversham, UK) built around an automated Nikon Eclipse Ti microscope equipped with a Perfect Focus System and an azimuthal TIRF unit (Gataca Systems, iLas2) using a 100X oil-immersion objective (Nikon CFI SR Apo, NA = 1.49), 488 nm and 640 nm simultaneous laser excitation at a TIRF angle of 80 deg, and two EMCCD cameras (Andor, iXon 888 Ultra) for fluorescent detection using a dichroic (Chroma, T565lpxr) to split the fluorescence onto the two cameras and a 525/50 bandpass filter (Chroma, 284337) for DAAM1-SNAP-Alexa488 detection and a 655 long-pass filter (Chroma, 283943) for SiR-actin detection. Imaging was performed at room temperature for 2 h taking 1 image every 2 min in 3 to 5 different sample locations chosen randomly and using 100 ms exposure time and 100 and 250 Electron Multiplying gain values, respectively for SiR-actin and DAAM1-SNAP-Atto488 detection.

TIRF-derived actin fluorescence micrographs were first denoised using rolling ball background subtraction (10 pixels), then contrast-enhanced using the Frangi Vesselness filter (Frangi et al. 1998). From the acquired time series, actin images were skeletonized and analyzed using publicly available plugins (Arganda-Carreras et al. 2010; Polder, Hovens, and Zweers, n.d.). The resulting skeleton maps were used as reference locations to quantify the quantity (intensity) of protein bound to the actin fibers. SNAP-tag protein fluorescence images were denoised using rolling ball background subtraction (50 pixels) and the pixel statistics were calculated with CLIJ ImageJ/Fiji plugins (Haase et al. 2020). All the scripts were written in ImageJ/Fiji (Schindelin et al. 2012), and are available upon request.

### Western Blot

Previously snap-frozen tissues or cell pellets were resuspended in RIPA buffer (150 mM NaCl, 1% Nonidet P-40, 1.0 mM EDTA, 1% Sodium Deoxycholate, 50mM Tris (pH 7.4)). The samples were subjected to sonication (1*10 sec pulse) and left on ice for 10 min. After centrifugation 5 min at 14000 rpm, the supernatants were collected and protein quantification determined using Bradford. Samples were resuspended in 4X SDS loading dye (200 mM Tris 6.8, 400 mM DTT, 4% SDS, 0.2% bromophenol blue, 20% glycerol), proteins were resolved by electrophoresis on a 10% SDS polyacrylamide gel and transferred on a nitrocellulose membrane. Blocking was performed in PBS 0.3% Tween, 5% milk. First antibody and secondary HRP labeled antibodies were diluted in the blocking buffer at a concentration provided in Table S2. Immunolabeling was detected by enhanced chemiluminescence and visualized with a digital luminescence image analyser Amersham Imager 600 (GE Healthcare).

### Co-immunoprecipitation Experiments

HA and 3xFlag tagged constructs were transiently transfected into 293T cells grown in 6 well plates using Lipofectamine 2000. After 48 hours, cells were harvested in cold phosphate-buffered saline (PBS), and pellets were resuspended in 250 μl of lysis buffer (50 mM Tris pH 7.4, 150 mM NaCl, 0.2% Triton X-100, 0.2% NP-40, and protease inhibitors). Lysates were cleared in a microcentrifuge by spinning at 15,000 g for 10 minutes at 4°C. Lysates were pre-cleared with 15 ul slurry of Dynabeads protein G (Thermo Fisher Scientific) with rotation at 4°C for 30 minutes. 5% of pre-cleared lysate was saved as input. 6 ug of anti-Flag M2 antibody (Sigma-Aldrich) was incubated with lysates for 1 hour at 4°C followed by incubation with 30 ul slurry of Dynabeads protein G washed in lysis buffer for 1 hour at 4°C with rotation. Following incubation, complexes were washed 5 times with lysis buffer. Elution was performed in 2.5x Laemmli buffer at 95°C for 5 minutes. The DAAM1 Co-IP signal was normalized to the IP signal of the FLAG-tagged protein being tested for interaction based on the intensity signal measured using ImageJ/Fiji software.

### Deletion of Daam1-MIC using CRISPR-Cas9 gene-editing

Mouse Embryonic Stem Cells (mESC, 129xC57Bl/6 background) were kindly provided by Kyung-Min Noh (EMBL Heidelberg) and cultured as described in Supplementary Methods. mESCs Knock-Out (KO) for the Daam1-MIC were generated using the CRISPR-Cas9 system (Ran et al. 2013) with a double guide RNA strategy (Sakuma et al. 2015). Each guide RNA targeted one of the two flanking introns and three gRNAs at each side were selected based on the proximity to Daam1-MIC and the quality score provided by (Doench et al. 2016). The best gRNA pair was chosen after testing all possible combinations and selecting the best editing efficiency. The primer sequences are provided in Table S1.

Gene editing was performed by transfection of 2 ug of Multiplex CRISPR vector ESCs with Lipofectamine 2000 (Invitrogen, 11668019). mESCs were plated at 2-3 different densities on 100 mm dishes (750 000, 1 500 000 and 3 000 000 cells/dish). Transfection of an empty vector was used as control. Six hours after transfection, the media was changed to prevent toxic effects. After 24h, puromycin selection was performed using a concentration of 1.5 ug/ml (Sigma, P8833), during 7 to 10 days. Afterwards, individual colonies were hand-picked into 96-well plates, expanded and genotyped by PCR and Sanger sequencing. Genotyping primer sequences are provided in Table S1. The confirmed KO clones and WT controls were further confirmed at the mRNA level using the RT-PCR primers amplifying Daam1-MIC described above.

### Immunofluorescence stainings and confocal imaging of cultured neurons

Neuronal differentiation from mESCs was done following the protocol reported by Bibel et al., 2007, with slight modifications (see Supplementary Methods) (Bibel et al. 2004; 2007). For immunofluorescence assays, cells on DIV0 or on DIV21 were washed with phosphate-buffered saline (PBS), fixed in 4% paraformaldehyde in PBS for 10 min, permeabilized in 0.3% Triton X-100 in PBS for 10 min, blocked for 1 h in 0.3% Triton X-100, % bovine serum albumin (BSA) in PBS and incubated in primary antibodies at 4oC overnight with shaking. Following this incubation, cells were incubated with the corresponding secondary antibodies (see Table S2) for 1 h at room temperature and mounted in FluoroShield with DAPI (Sigma, F6057-20ml) for imaging. Images were taken on an SP8 confocal microscope (CRG, Advanced Light Microscopy Unit) using identical settings for each condition in a given experiment. Briefly: for IEG positive nuclei analysis (Figure; DIV21) and neurite length analysis (Figure ; DIV0+4h’s) dry 20X objective was used to image the whole span of the neuronal culture (17-23µm, Z-step size 1µm, zoom factor 2.5). For filopodia analysis 63X oil-immersion objective was used to image the whole span of the protrusion (around 2 µm, Z-step size 0.12 µm, zoom factor 2.5). Confocal sections were Z stack projected with maximum intensity selection and analyzed in ImageJ/Fiji software. Immediate early genes positive nuclei were counted using Analyze Particles plugin (nuclei size at least 25 pixel), and normalized to DAPI positive nuclei. Neurite and filopodia length were analyzed manually using the Segmented *Line* plugin.

### Measurement of intracellular [Ca2+] in cultured neurons

Cytosolic Ca2+ signal was determined at room temperature in cells loaded with 4,5 mM FURA-2AM (30 min incubation at 37ºC). Ca2+ responses were calculated as the ratio of emitted fluorescence (510 nm) after excitation at 340 and 380 nm, relative to the ratio measured before cell stimulation (FURA-2AM ratio 340/380). Briefly, cells were maintained in an isotonic solution containing (in mM): 140 NaCl, 5 KCl, 1.2 CaCl2, 0.5 MgCl2, 5 glucose, 10 HEPES (305 mosmol/l, pH 7.4 with Tris) for 2 minutes and then stimulated with a 30 mM KCl isotonic solution (115 NaCl, 30 KCl, 1.2 CaCl2, 0.5 MgCl2, 5 glucose, 10 HEPES) to activate voltage-gated calcium entry. As indicated in the respective figure legends, cells were treated with 5 µM Latrunculin A (LatA), 50 µM small-molecule inhibitor of Formin Homology 2 domains (SMIFH2) or vehicle (DMSO). The treatment was maintained for the experiment (40 minutes). In the case of SMIFH2, its effect on neuronal cultures was measured after 90 min exposure to 50 µM SMIFH2. Y-27632 ROCK inhibitor was used at a concentration of 10 µM, and the effect was measured immediately. All experiments were conducted at room temperature as described previously (Fernandes et al. 2008). AquaCosmos software (Hamamatsu Photonics) was used for capturing the fluorescence ratio at 510 nm obtained post-excitation at 340 and 380 nm, respectively. Images were computed every 5 sec. Measurements were processed using SigmaPlot 10 software.

### Generation of mouse Daam1-MIC KO mice

Chimeric mice were obtained by blastocyst injection of one of the Daam1-MIC KO mESC lines into B6 albino (B6(Cg)-Tyrc-2J/J) embryos, which were then transferred to pseudopregnant CD1 females. Chimeric males were subsequently crossed to albino B6 females, and all non-albino mice were genotyped to select Daam1-MIC KO mice. These were backcrossed four times to C57Bl/6J and heterozygous mice were then crossed to generate Daam1-MIC KO and wild type (WT) littermates for experiments. The colony was maintained at the Animal Facility of the Barcelona Biomedical Research Park (PRBB). All procedures were approved by the PRBB Animal Research Ethics Committee and the Generalitat de Catalunya and were carried out in accordance with the guidelines of the European Union Council (2003/65/CE) and Spanish regulations (BOE 252/34367-91, 2005). WT mice were purchased from Charles River (references: 709 for albino B6, 022 for CD1 and 632 for C57Bl/6).

### Functional synapse quantification in mice hippocampus

7 control mice and 8 DAAM1-MIC mice of 22 days postnatal (PD22) were perfused 5 minutes with 4% paraformaldehyde and brains were transferred to Phosphate Buffered Saline immediately. Brains were cut in 70µm sagittal slices, and equivalent slices between animals were selected for immunohistochemistry. Slices were placed in a blocking/permeabilization solution containing 5% horse serum and 0.25% Triton X-100 in PBS for 1 hour, followed by immunostaining with primary antibodies (Table S2) two days at 4°C. Slices were washed 4-5 times in PBS before staining with secondary antibodies (Table S2) overnight at 4°C. Slices were labeled with 4′,6-diamidino-2-phenylindole (DAPI) (Sigma-Aldrich, D9542) for 5 minutes and washed 3-4 times in PBS before being mounted in Mowiol mounting media (Millipore, 475904-M).

CA1 hippocampus was imaged with a 63x oil immersion objective on a Leica SPEII confocal microscope. ROIs with a z-stack of 5µm and a z-step size of 0.2µm were captured. Three ROIs per slice, and three slices per animal were analyzed with Imaris software. Images were processed with Background subtraction and Median filter for both channels, followed by segmentation with Imaris models. To quantify the number of functional synapses the “Spots Close To Surface XTension” of MatLab was applied with a threshold of 0.5µm.

DAAM1-MIC results were normalized to control mean and unpaired Mann-Whitney nonparametric test was performed for statistical analysis.

### Immunohistochemical imaging of mice hippocampus

Mouse hippocampal sections were prepared as previously described (Hoeymissen et al. 2020). In brief, mice were euthanized with CO2 and perfused transcardially with 0.1 M PBS, followed by 4% paraformaldehyde in PBS until tissues were completely cleared of blood. Fixed brains were extracted and stored in 4% paraformaldehyde at 4 °C for 24 h, and in sucrose 30% with 0.01% azide in PBS for the following 24 h. Prepared tissues were cut into 40 µm coronal sections, in serial order throughout the dorsal hippocampus (Bregma sections between -1.34mm to -2.54mm). Sectioning was performed by an in-house Tissue Engineering Unit (CRG). Immunohistochemistry was done by tissue permeabilization in 0.5% Triton X-100 in PBS for 15 min × 3 times, blocked for 2 h in 10% normal goat serum (NGS) in PBS albumin and incubated primary antibodies in 5% NGS, Tween 0.5% in PBS at 4oC overnight. Following this incubation, cells were incubated with the corresponding secondary antibodies (see Table S2) for 2 h at room temperature. Washing steps were repeated (for 15 min × 3 times) and sections were mounted in FluoroShield with DAPI for imaging. Images were taken on a confocal microscope (SP8 Leica; CRG, Advanced Light Microscopy Unit) using identical settings for each condition in a given experiment with a dry 20X objective. A single middle plane of each section was imaged and the images were analyzed in ImageJ/Fiji software. DG size was marked manually with Polygon selections using DAPI stained nuclei as a region of reference marker and consecutively quantified with the Measure plugin. A detailed list of all stainings and primary antibodies used is provided in Table S2.

### Behavioral and locomotor tests in a neonatal mice

A battery of behavioral and motor tests to probe early post-natal neurodevelopment was performed as described in Feather-Schussler and Ferguson (2016) and Roper et al. 2021, with some adjustments, as detailed in Supplementary Methods. In particular, we performed the following tests at PDs 4, 7, 10 and 14, unless stated otherwise: (i) Pivoting and walking, (ii) Righting reflex, (iii) Preyer’s reflex, (iv) Front-limb suspension, (v) Hindlimb suspension, (vi) Grasping reflex, (vii) Cliff aversion, (viii) Negative geotaxis, (ix) Homing test.

### Behavioral tests in Adult Mouse

We also performed a battery of behavioral and locomotors tests in adult mice. Each test was performed twice, using six animals per sex and genotype in each replicate. All mice were between 2 and 5 months old and siblings were matched when possible. As described in detailed in Supplementary Methods, we performed the following tests: *(i) Spontaneous basal locomotor activity, (ii) Novel Object Recognition (NOR), (iii) Elevated Plus Maze, (iv) Morris Water Maze, (v) Grip strength, (vi) Rotarod, (vi) Beam Balance, (vii) Fear conditioning*.

### Rescue experiments with ROCK inhibitor in Adult Mice

The ROCK inhibitor (Y-27632; HelloBio HB2297) trans-4-[(1R)-1-aminoethyl]-N-4-pyr-idinylcyclohexanecarboxamide was dissolved in physiological saline at a concentration of 1mg/ml and injected for three consecutive days intraperitoneally at a dose of 10mg/kg (Li et al. 2009). The equivalent volume of saline solution was injected into the control mice. On the 7th day post-injection, NOR experiments were performed as described in Supplementary Methods.

### Electrophysiology ex vivo

Mice (8–16 week old) were decapitated immediately, the brain was quickly removed and submerged in artificial cerebrospinal fluid rich in aCSF sucrose buffer (2 mM KCl, 1.25 mM NaH2PO4-H2O, 7 mM MgSO4, 26 mM NaHCO3, 0.5 mM CaCl2, 10 mM glucose and 219 mM sucrose) at 4°C, saturated with a 95% O2, 5% CO2 mixture and maintained at pH 7.32-7.4. Transverse brain slices (300 µm thick) were cut with a vibratome (VT1200S, Leica) in oxygenated aCSF sucrose at 4°C and transferred to recovery chamber with oxygenated aCSF buffer 124 mM NaCl, 2.5 mM KCl, 1.25 mM NaH2PO4-H2O, 1 mM MgSO4, 26 mM NaHCO3, 2 mM CaCl2 and 10 mM glucose and incubated for > 1 h at room temperature (21-24 ºC) and pH 7.32-7.4. Individual slices were transferred to an immersion recording chamber and perfused with oxygenated aCSF at 2 mL/min (30 ± 2°C).

Whole-cell intracellular recordings in voltage clamp (VC) and current clamp (CC) mode were performed in pyramidal neurons in CA1 stratum pyramidale. Cells were visualized with a water-immersion 40x objective. Patch electrodes were fabricated from borosilicate glass capillaries (P1000, Sutter Instrument) with resistance of 4–6 MΩ when filled with the internal solution that contained : 130 mM K-MeSO4, 10 mM HEPES, 0.5 mM EGTA, 2 mM MgCl2, 4 mM Mg-ATP, 0.4 mM Na-GTP, 10 mM phosphocreatin di (tris) salt and 0.3 % biocytin, for membrane properties experiments in CC and excitatory postsynaptic currents (EPSC) evoked in VC, in pyramidal neurons; 130 mM Cs-MeSO4, 5 mM CsCl, 10 mM HEPES, 0.5 mM EGTA, 2 mM MgCl2, 4 mM Mg-ATP, 0.4 mM Na-GTP, 10 mM phosphocreatin di (tris) salt and 0.3 % biocytin, for miniature excitatory postsynaptic currents (mEPSC) in VC recordings from pyramidal neurons. All pipette solutions were adjusted to pH 7.2-7.3 with K-OH or Cs-OH. Membrane currents and voltages were acquired with Multiclamp 700B amplifiers, digitized (Digidata 1550B) and controlled by pClamp 10.7 (Molecular Devices Corporation, Sunnyvale, CA, USA) software. Membrane intrinsic properties of CA1 pyramidal cells were determined by passing hyperpolarizing and depolarizing current steps (1 s, with 10 pA increments from -100 to 140 pA) in CC.

Synaptic responses in CA1 were evoked by monophasic current (50 μs duration) stimulation of the Schaffer collateral fibers (SCs) with an extracellular bipolar tungsten electrode via isolated current stimulator (DS3) that was set to deliver monophasic currents of 50 μs duration. Membrane currents and voltages were acquired with Multiclamp 700B amplifiers, digitized (Digidata 1550B) and controlled by pClamp 10.7 (Molecular Devices Corporation, Sunnyvale, CA, USA) software. To study changes in the probability of transmitter release of the presynaptic cell (e.g., Bekkers and Stevens 1990; Kullmann 1994; Malinow and Tsien 1990) we applied paired-pulse facilitation (PPF) protocol, which consisted of evoking two consecutive EPSC responses at different intervals (from 20 ms at 160 ms, a 20 ms inter-pulse interval). Changes in the PPF were calculated as a PPF ratio from (R2-R1)/R1, where R1 and R2 were the peak amplitudes of the first and second EPSCs, respectively (Martin and Buño, 2002). Extracellular field postsynaptic potentials (fEPSPs) were recorded by placing borosilicate glass electrode filled with aSCF sucrose buffer in the stratum radiatum (SR) of the CA1 pyramidal layer. Evoked fEPSPs were elicited by stimulation of the SCs fibers as EPSCs.

For LTP experiments, the stimulus intensity was adjusted to elicit 50% of the maximum response signal and kept constant throughout the experiment. Data was stored through an acquisition system (Axon Instruments) and the software pClamp 10.7 was used to display fEPSP and measurements of fEPSP slopes. After recording stable baseline responses for 30 min, LTP was induced by a single train of theta burst stimulation (TBS; 10 bursts of 5 pulses at 100 Hz, with an interval of 200 ms between bursts). Potentiation was measured for 1 h after LTP induction at 0.033 Hz. Changes in the fEPSP slope were calculated in relation to the baseline fEPSP responses during the last 10 min before TBS, and the time course of LTP values was then normalized to this baseline.

### Dendritic spine morphology analysis

For spine analysis, neurons were filled with biocytin (Sigma-B4261) during whole-cell recordings. Slices (300 µm) with recorded cells were fixed overnight with 4% paraformaldehyde in PBS at 4°C and then transferred to 0.05 % Na-azide in PBS. After slices were permeabilized and blocked with 0.3% Triton X-100 in PBS and 10% normal goat serum (NGS) for 1 h at room temperature. To reveal biocytin, the slices were incubated overnight at 4°C with streptavidin Alexa Fluor-488 conjugated (1:1 000) in 0.3% Triton X-100 in PBS. Confocal microscopy (Leica SP8; CRG, Advanced Light Microscopy Unit) was used to capture fluorescence Z-stacks of dendritic spines from biocytin filled cells with a 0.2 µm step using a 63X glycerol immersion objective. Only dendrites with bright and continuous labeling were included for analysis. Primary dendrites were not included in the analysis. Huygens’ essential software was used to deconvolve images. All images were batch processed using the same template (available upon request). Deconvoluted images were imported into NeuronStudio and semi-automatically analyzed blind to experimental conditions. Dendritic spine quantification was performed on a total of 23 CA1 pyramidal cells (n = 11-12 cells per group; n = 2 groups). Spine density was measured at least in 100 µm of secondary apical branches. All dendritic spines were at least 50 µm apart from the neuronal soma. The total number of spines was divided by the total length of the dendritic spines.

### RhoA2G FRET biosensor analysis of neuronal cells by microscopy

To obtain lentivirus for infection of neurons, HEK293T cells were seeded in lentivirus packaging media (94.99% Opti-MEM I, 5% FBS,0.01% Sodium Pyruvate) were transfected according to manufacturer instructions using Lipofectamine 2000 with plasmids carrying HIV-1 Gag/Pol (pMDLg/pRRE Addgene: 12251), HIV-1 Rev (pRSV-Rev Addgene: 12253), and VSV g-glycoprotein Env (pMD2.G Addgene: 12259), together with pLentiRhoA2G (Addgene: 40179). Media was changed after 6 h to neuronal N2 media. The viral particles were harvested from the neuronal N2 media 48 h after the transfection and used for cell infection. Experiment was performed on two separate neuronal differentiations considered as replicates. Each differentiation was performed using three KO and three WT cell lines (described before), and fixed DIV21 neurons were used for FRET analysis. Procedure was performed in accordance with safety rules and approved by the CRG Biosafety committee (procedure registration number CBS19_015_A).

Relative FRET was calculated automatically using AcceptorPhotobleaching (confocal microscope SP8 Leica; CRG, Advanced Light Microscopy Unit) by dividing the normalized fluorescence intensity of Venus at its emission peak (528 nm) by the normalized fluorescence intensity of mTFP1 at its emission peak (492 nm). Provided Lookup Table (LUT) is linear and covers the full range of the data.

### RNA-sequencing and transcriptomic analyses

In vitro differentiated glutamatergic neurons (DIV21) were exposed to 10 µM Y-27632 ROCK inhibitor or vehicle (water) for 2 hours, in triplicates. Next, the cells were collected by scraping and snap-frozen in liquid nitrogen. These samples were subjected to RNA extraction using the RNeasy Plus Mini kit (Qiagen, 74136). Library preparation (polyA-selected, stranded) and RNA sequencing were performed by the CRG Genomics Unit following the standard Illumina protocol. An average of 35.3 million 50 bp single-end reads were generated for each library on a NextSeq 2000 machine. RNA-seq data was deposited in Gene Expression Omnibus (GEO), and is available under the GSE219244 accession number.

Reads were aligned to the Mus musculus transcriptome assembly from Ensembl (release 104; GRCm39) with Salmon v.1.4.0 (Patro et al. 2017). Raw gene counts and TPMs were extracted from Salmon outputs using the tximport (Soneson, Love, and Robinson 2015). To identify mis-regulated genes in the KO that were “rescued” upon ROCK inhibition, we performed the following steps. First, differential gene expression between Daam1-MIC KO and WT samples independently of the treatment was calculated using DESeq2 on the raw reads (Love, Huber, and Anders 2014) with a model for genotype and treatment; p-values were corrected using the Independent Hypothesis Weighting (IHW) (Ignatiadis et al. 2016). Second, we again used DESeq2 to independently compare gene expression profiles from each condition against the control KO one (i.e., KO exposed to water [Veh]). Raw p-values from Wald Test from the three independent comparisons (WT-Veh vs KO-Veh, WT-ROCKi vs KO-Veh, and KO-ROCKi vs KO-Veh) were combined into a single p-value with the Stouffer method, weighting double the p-values from the comparison between inhibitor and vehicle in KO samples (KO-ROCKi vs KO-Veh), to emphasize the response of the KO samples to the inhibitor on gene expression. Next, we labeled the genes differentially expressed independently of the treatment as “non-rescued” and removed them from subsequent analyses. P-values of the remaining genes were then corrected with IWH (Ignatiadis et al. 2016) using the mean gene TPMs as covariates. Genes with a combined adjusted p-value lower than 0.01 were then considered to present a rescue pattern since their expression differs in the control KO without the inhibitor (KO-Veh) compared to the rest of the conditions. Gene Ontology term enrichment for rescued genes was calculated using the Bioconductor Clusterprofiler package (Wu et al. 2021). Lastly, we used pathview (Luo and Brouwer 2013) to visualize log2 fold changes on genes in KEGG pathways.

## End Matter

### Author Contributions and Notes

P.P performed molecular, cellular, and behavioral characterization of the microexon function, with the help of M.M.C. A.Z.M generated and analyzed electrophysiology data (Figure 6 and S8); F.M. performed the analysis of the actin polymerization assay, provided support with bioinformatic analyses and contributed with critical insights. G.C.R performed calcium imaging experiments and the analysis. D.N. performed TIRF and dual-color TIRF microscopy experiments and R.G.R. wrote ImageJ/Fiji scripts necessary for the downstream analysis. L.P.I. analyzed RNA-seq data and provided help with animal behavior data analysis. C.M.P and P.O.C. performed functional synapse quantification (Figure 4) . S.B. performed a Western Blot analysis and contributed with critical insights. C.R.-M. and J.P. performed RT-PCR assays and genotyping. M.M. De L.C., Á.F.-B., C.S., J.L.M.L., and O.F. supported the animal experiments performed. D.L. and J.I.W.F. helped with FRET acceptor photobleaching experiments. J.E. performed Co-IP experiments, supervised by B.B. P.P., E.H., M.D., and M.I., and designed the experiments with input from other authors. P.P., M.D., and M.I. conceived the study, supervised the work, and wrote the manuscript with input from other authors.

The authors declare no conflict of interest.

## Acknowledgments

We thank Kyung-Min Noh for providing the mESC line, Miguel Valverde and Francisco Muñoz for their help and feedback with the calcium imaging experiments, and members of the Irimia and Dierssen groups for constant scientific discussion and feedback. We also thank the CRG Genomics, Protein Technologies, Tissue Engineering and Advanced Light Microscopy Units for the RNA sequencing, protein purification, blastocyst injection and microscopy services, respectively.

## Supplementary figures

**Supplementary Figure 1.**
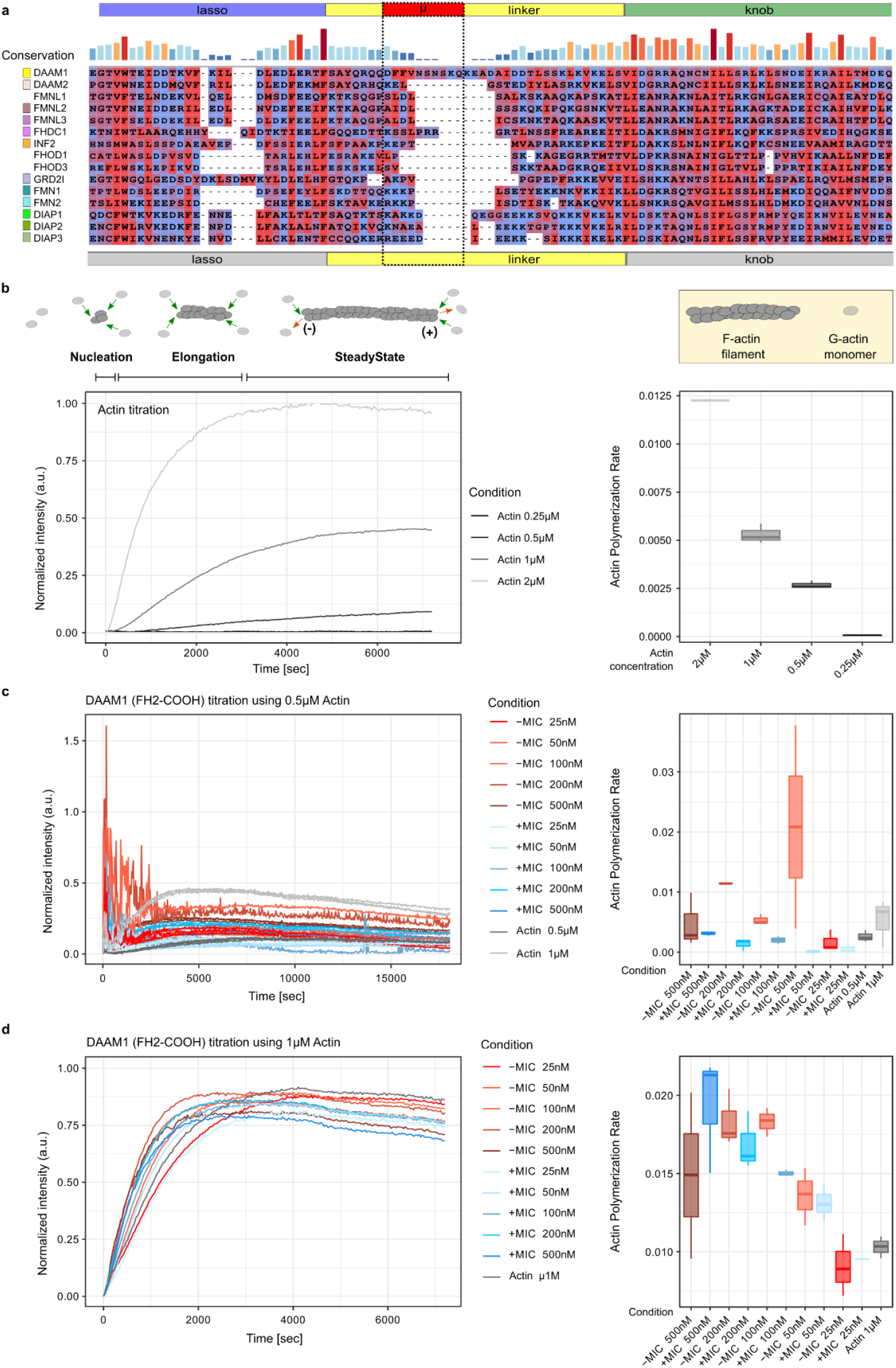
Linker region conservation among formins and actin pyrene assay optimization. (a) Amino acid sequence alignment of the linker regions of formin proteins. (b) Concentration-dependent actin self-assembly. Left: actin self-assembly measured by incubating the pyrene-labeled actin at various concentrations. Top scheme: schematic representation of nucleation, elongation, and steady-state phase of actin filament assembly based on 1 μM actin curve. Green arrows describe the potential incorporation of G-actin monomer into F-actin filament while red arrows indicate dissociation. Right: distribution of actin polymerization rates per condition calculated according to Doolittle et al. 2013. (c,d) Left: actin self-assembly activity at various concentrations of the two splice variants of the DAAM1 FH2-COOH fragment using 0.5 μM (c) and 1 μM actin (d). Actin assembly was measured by incubation of the pyrene-labeled actin. Right: distribution of actin polymerization rates per condition calculated based on Doolittle et al. 2013.

**Supplementary Figure 2.**
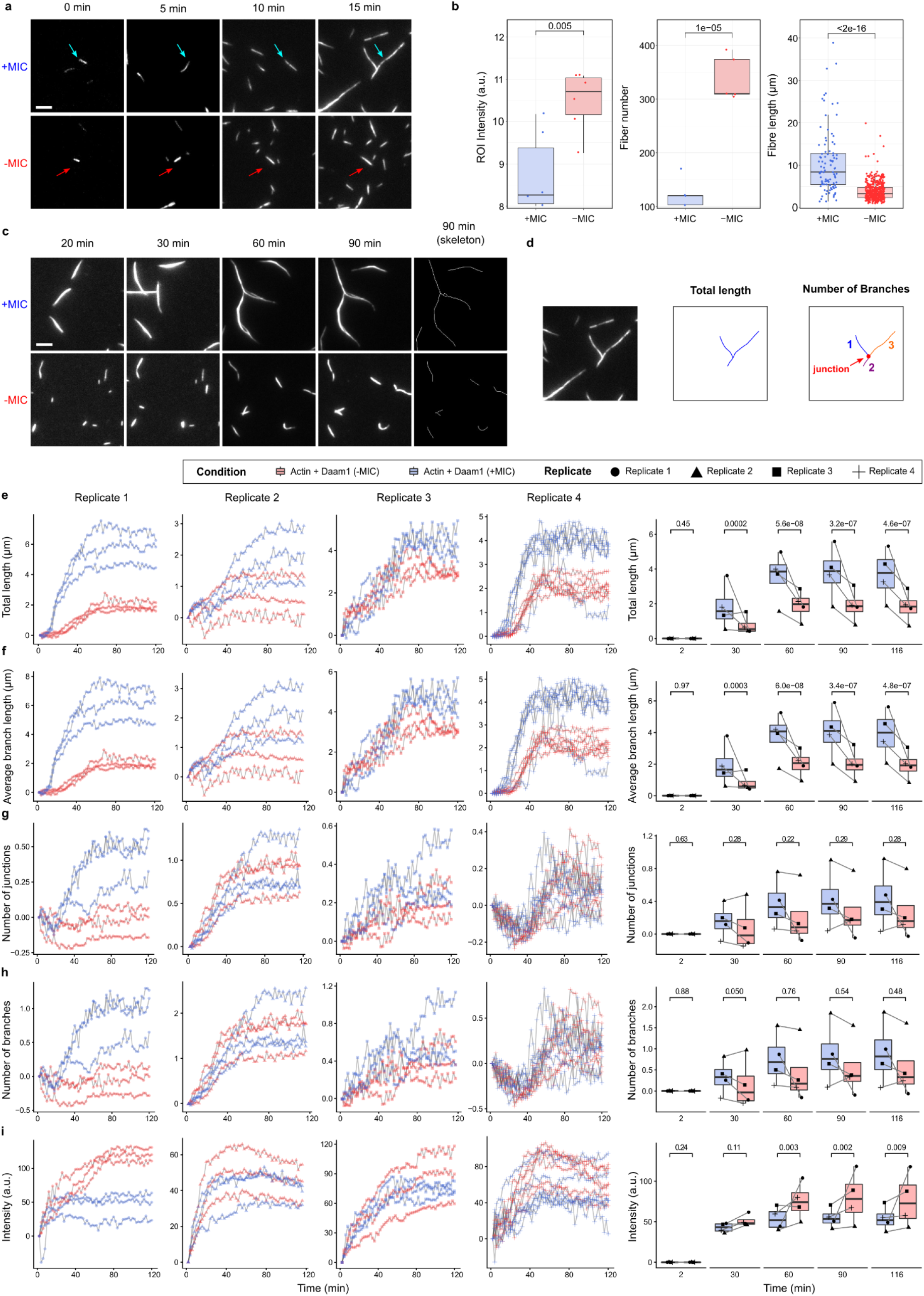
TIRF microscopy of actin polymerization. (a) Magnification of the actin fluorescence TIRF micrographs shown in Figure 2g is used for tracking the behavior of individual actin fibers over time (examples indicated by the colored arrows). Scale bar: 2 μm. (b) Quantifications of ROI intensity, fiber number, and length from (Figure 2g) at the last recorded time point (15 min). Each dot (in the left and central box plot) corresponds to individual ROIs that were randomly chosen from the sample. P-values from Wilcoxon rank-sum tests. (c) Magnification of actin fluorescence TIRF micrographs obtained using 0.2 μM actin and 200nM FH2-COOH Daam1 fragments. The right panels show the skeletonized actin fibers (90 min skeleton) as obtained using the AnalyzeSkeleton Fiji plugin (Schindelin et al. 2012; Polder et al. 2010). Scale bar: 2 μm. (d) Schematic representation of the morphological features quantified in (e-h). (e-i) Temporal quantifications of F-actin fiber features including fiber total length in μm (e), average length of individual branches in μm (f), number of junctions per object (g), number of branches per object (h), and actin fiber intensity (i). Results from three different fields of view are shown per each replicate and per variant (blue: +MIC, red: -MIC). Right: Boxplots summarizing the average values of the quantified F-actin fiber features. Black lines in the box plots connect data points acquired simultaneously during the course of a single experiment. P-values from two-way ANOVA tests with replicates and as factors.

**Supplementary Figure 3.**
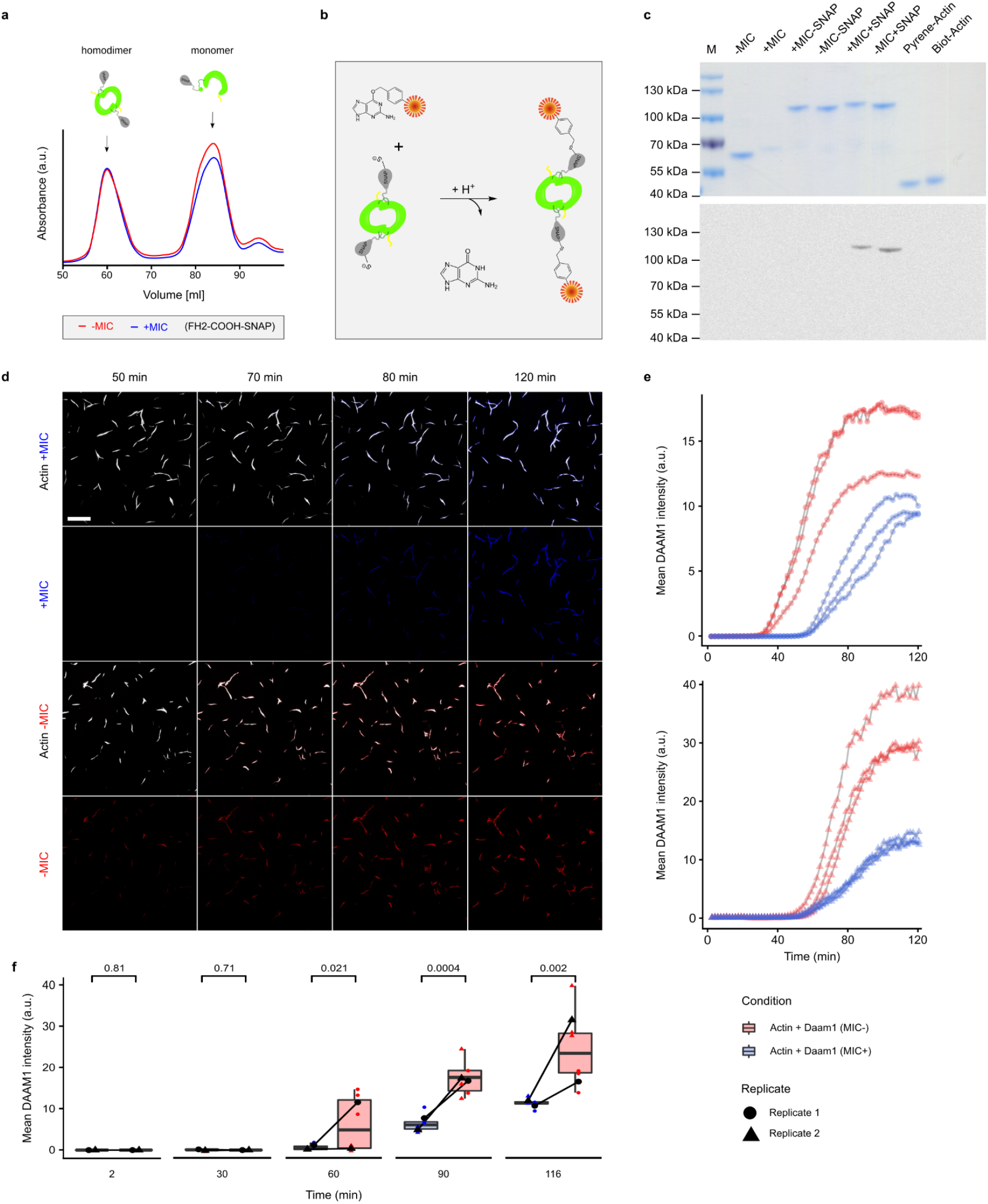
Dual-color TIRF microscopy with SNAP labeling. (a) FPLC elution profiles of DAAM1 SNAP-FH2-COOH fragments purified from bacteria. FPLC was performed using a HiLoad® 16/600 Superdex® 200 pg column. (b) Schematic representation of fluorescence labeling of purified SNAP-FH2-COOH fragments. Green indicates the DAAM1 fragment, gray is the SNAP-tag, and orange is the fluorescent labeling of the SNAP-tag with Alexa Fluor 488. (c) Top: Coomassie blue-stained SDS-PAGE gel showing all the proteins used throughout the course of this study. Bottom: A fluorescence image of the SDS-PAGE gel is shown in the top panel using blue light excitation. (d) Representative micrographs of dual-color TIRF experiments performed using 0.2 μM actin and 200 nM SNAP-tagged proteins. Scale bar: 10 μm. (e) Quantification of the mean protein fluorescence intensity per actin fiber throughout the course of the experiment in Panel d for replicate 1 (upper panel) and replicate 2 (lower panel). (f) Quantification summary for both replicates from Panel e. Data plots show multiple regions analyzed per each experimental replicate and boxplots summarize their averages. Gray lines connect data points acquired simultaneously during the course of a single experiment. P-values from two-way ANOVA tests with replicates and as factors.

**Supplementary Figure 4.**
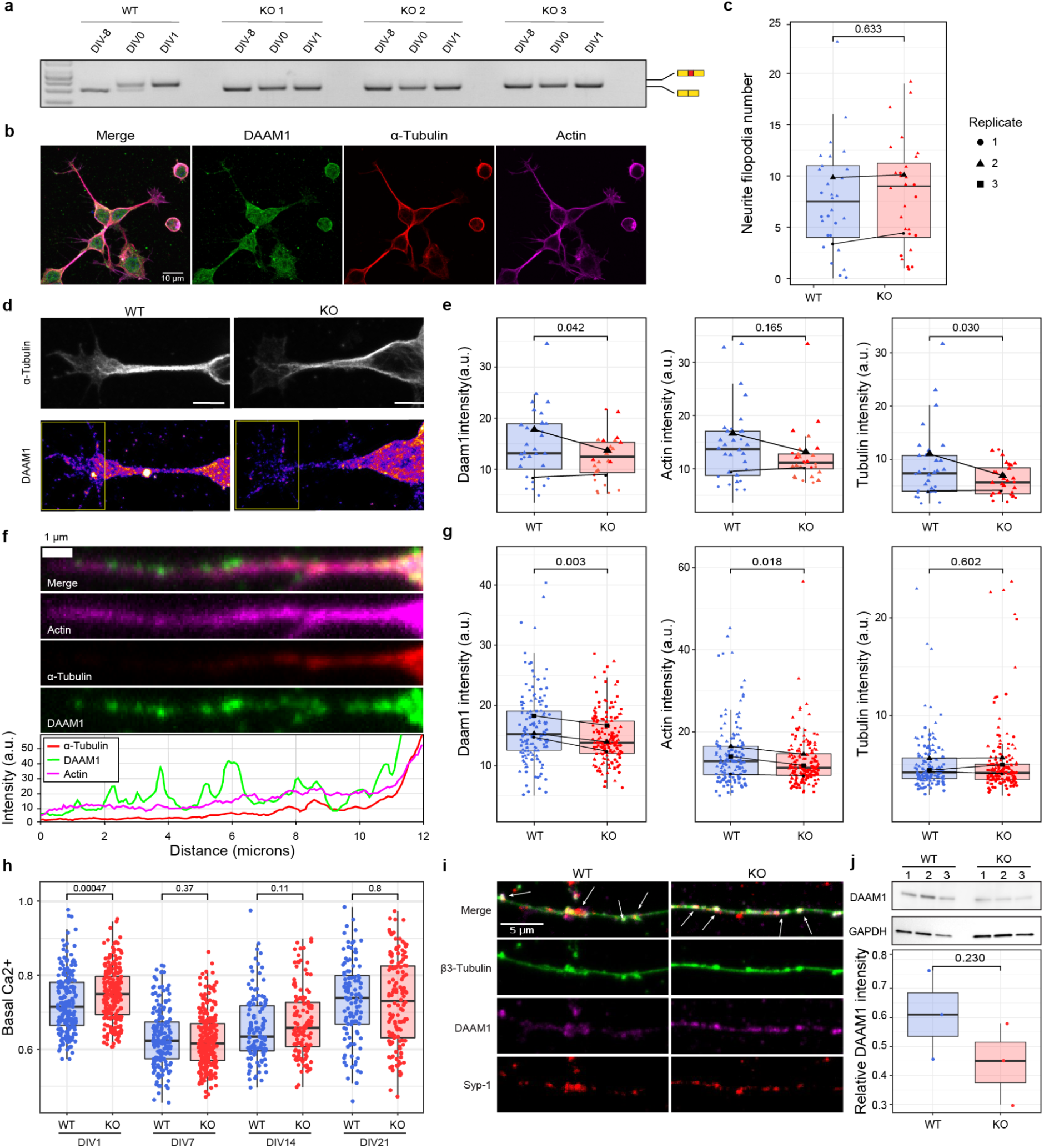
Increased filopodia and neurite length upon microexon removal in neuronal precursors. (a) RT-PCR assays of Daam1-MIC inclusion during neuronal differentiation in WT and KO cell lines. (b) Representative immunocytochemistry image of WT neuronal precursors (NPCs; DIV0+4h). Scale bar: 10 μm. (c) Distributions of neurite filopodia number in NPCs (DIV0+4h). Dots represent values measured on an individual neurite. (d) Representative immunocytochemistry images of derived growth cones from WT and KO NPCs (DIV0+4 h). Relative intensity throughout the last 5 μm of the growth cone (yellow boxes) was analyzed. (e) Distributions of relative intensities of DAAM1, actin, and tubulin. Dots represent individual values per growth cone. (f) Top: representative immunocytochemistry images of a filopodia from a WT NPC (DIV0+4 h). Bottom: Intensity spectra throughout the filopodia. (g) Distributions of relative intensities of DAAM1, actin, and tubulin. Dots represent values measured on the last 5 μm from the tip of individual filopodia. (h) Basal calcium flux comparison between WT and KO cell lines, based on FURA 2 AM ratio throughout the time course of the neuronal differentiation protocol. (i) Representative immunocytochemistry image of a DIV21 neuronal protrusion stained with β3-Tubulin, DAAM1 and synaptic marker Syp-1. Arrows highlight the overlap between DAAM1 and Syp-1 puncta. (j) Western blot (left) and associated quantification (right) of the relative intensity of DAAM1 normalized to GAPDH. Protein extract from neuronal cultures DIV21. Dots represent values measured on individual cell lines. P-values from Wilcoxon rank-sum tests (h), two-way ANOVA tests with replicate and genotype as factors (c,e,g), or Student’s t-test (j).

**Supplementary Figure 5.**
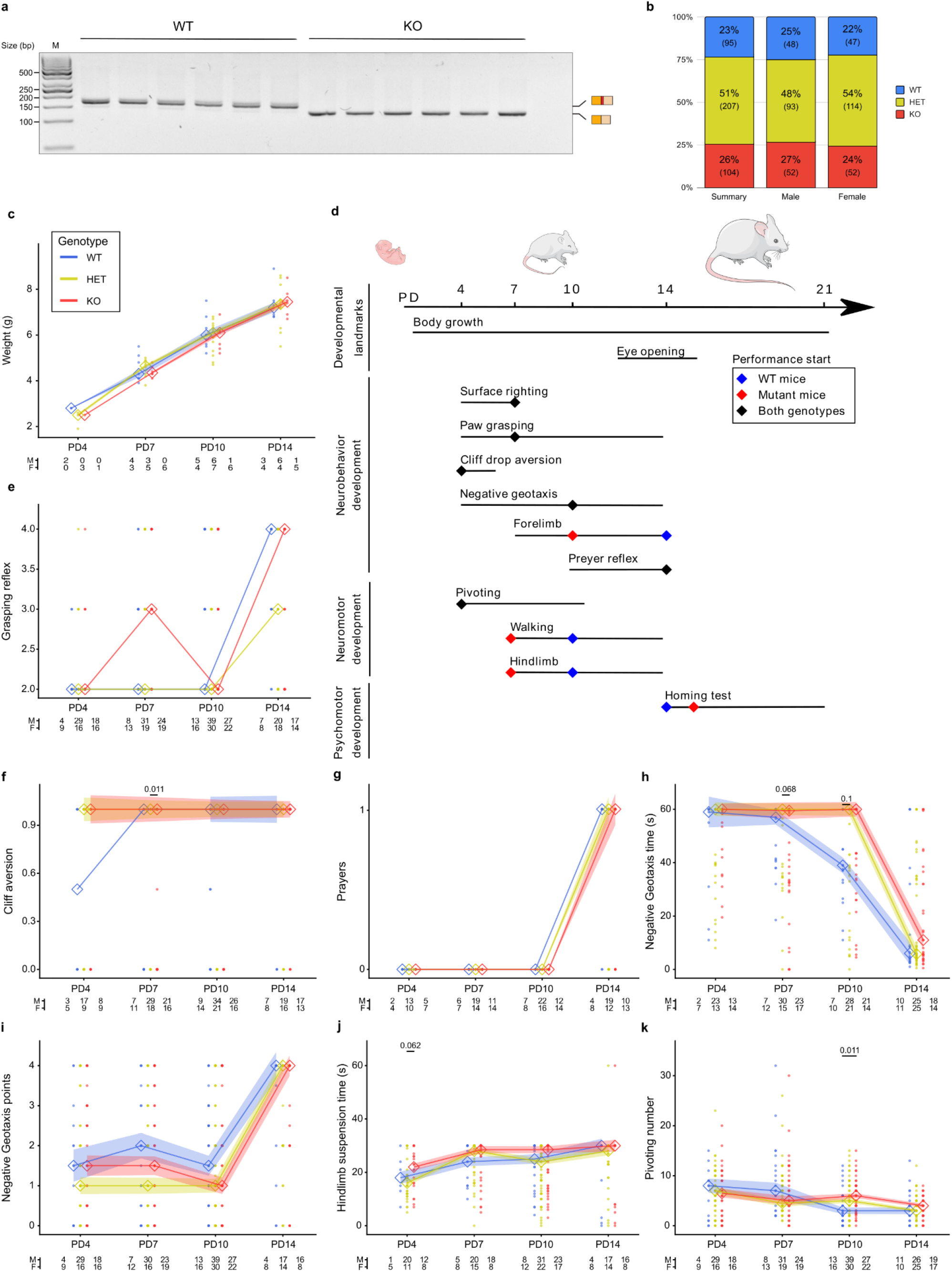
Results from various neonatal experiments. (a) RT-PCR assays of Daam1-MIC inclusion in WT and KO mice at postnatal day (PD) 21. (b) Distribution of Mendelian ratios for all mice used in this study (N = 406). Numbers in brackets represent the animal number. (c) Lineplot representation of the animal weight. (d) Mouse postnatal developmental timeline and summary of the results of the neurodevelopmental experiments performed on days 4, 7, 10, 14, and 21. Lines represent the span between the start and the end of each experiment, while diamonds indicate when >50% of animals of a given genotype achieve a specific level of performance. Blue and red diamonds correspond to WT and mutant mice, respectively, whereas black diamonds indicate similar performance. PD - postnatal days. Mouse icons are derived from bioicons.com. (e-k) Lineplot representation of the animal performance in various tests during the experimental time course (see Supplementary Methods for details). Thick lines represent the mean performance for both sexes, shading represents the standard error mean (SEM). The bottom numbers represent the number of males (M) and females (F) used. Dots represent values measured per animal. P-values from Wilcoxon rank-sum tests against the WT.

**Supplementary Figure 6.**
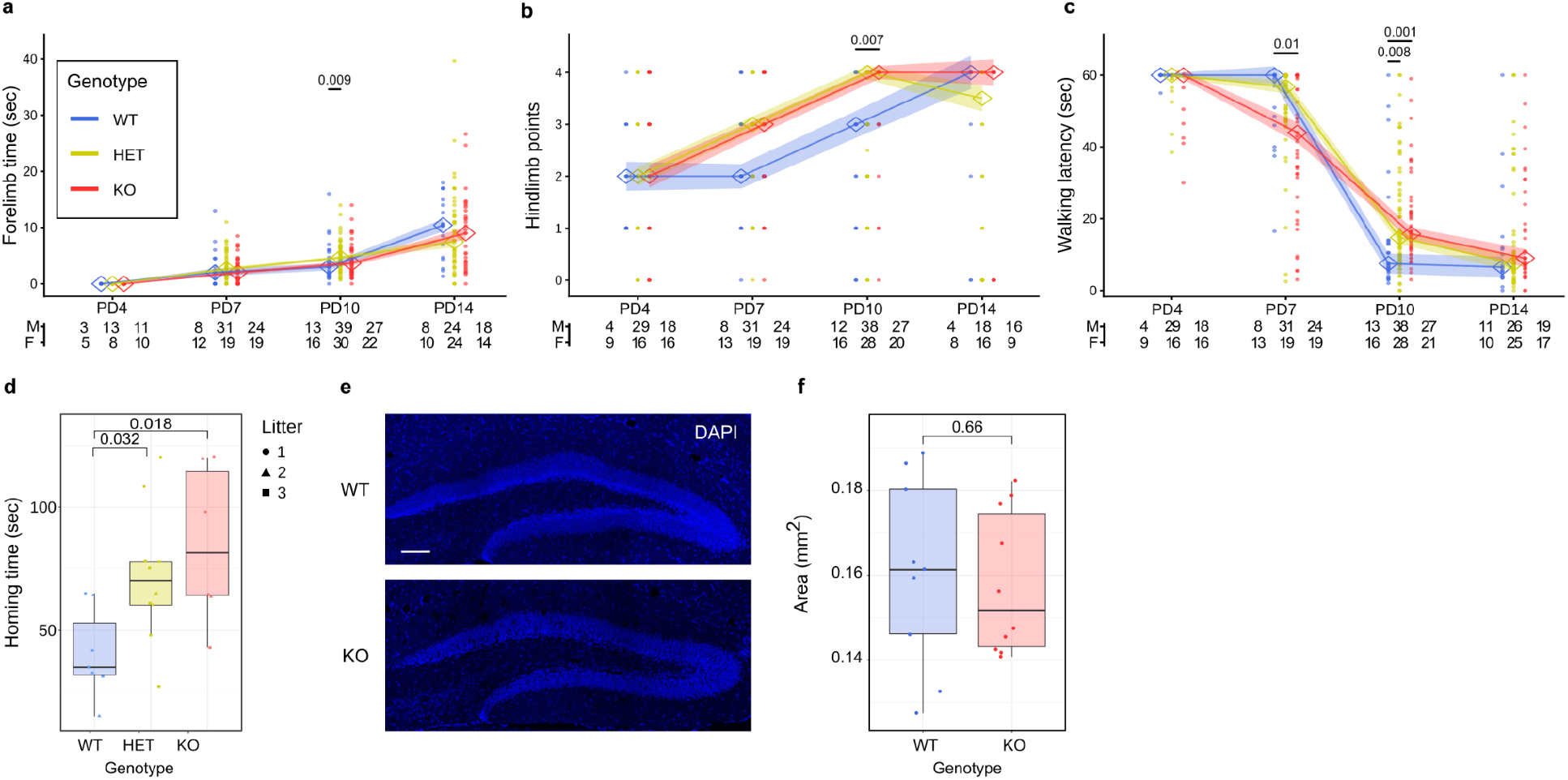
Results from various neonatal experiments. (a-c) Animal performance at each indicated time point for forelimb strength (a), hindlimb suspension score (b), and latency to walk (c). Thick lines represent mean performance, shading represents the standard error mean (SEM). Dots represent values measured per animal. The number of animals per time point, genotype (WT, heterozygous [HET] and homozygous [KO] mutant), and sex (males [M] and females [F]) are shown at the bottom. P-values from Wilcoxon rank-sum tests against the corresponding WT. (d) Latency to reach the target area in the homing test per genotype. Values for matched litters are indicated with symbols. P-values from Wilcoxon rank-sum tests. (e) Representative images of DAPI stained nuclei (blue) in sections of the dentate gyrus of the hippocampus from PD21 mice (PD21). (f) Quantification of dentate gyrus size across images. One dot represents one animal, where an average of 3 to 6 sections of coronal view of the hippocampus were analyzed. P-values from Wilcoxon rank-sum tests.

**Supplementary Figure 7.**
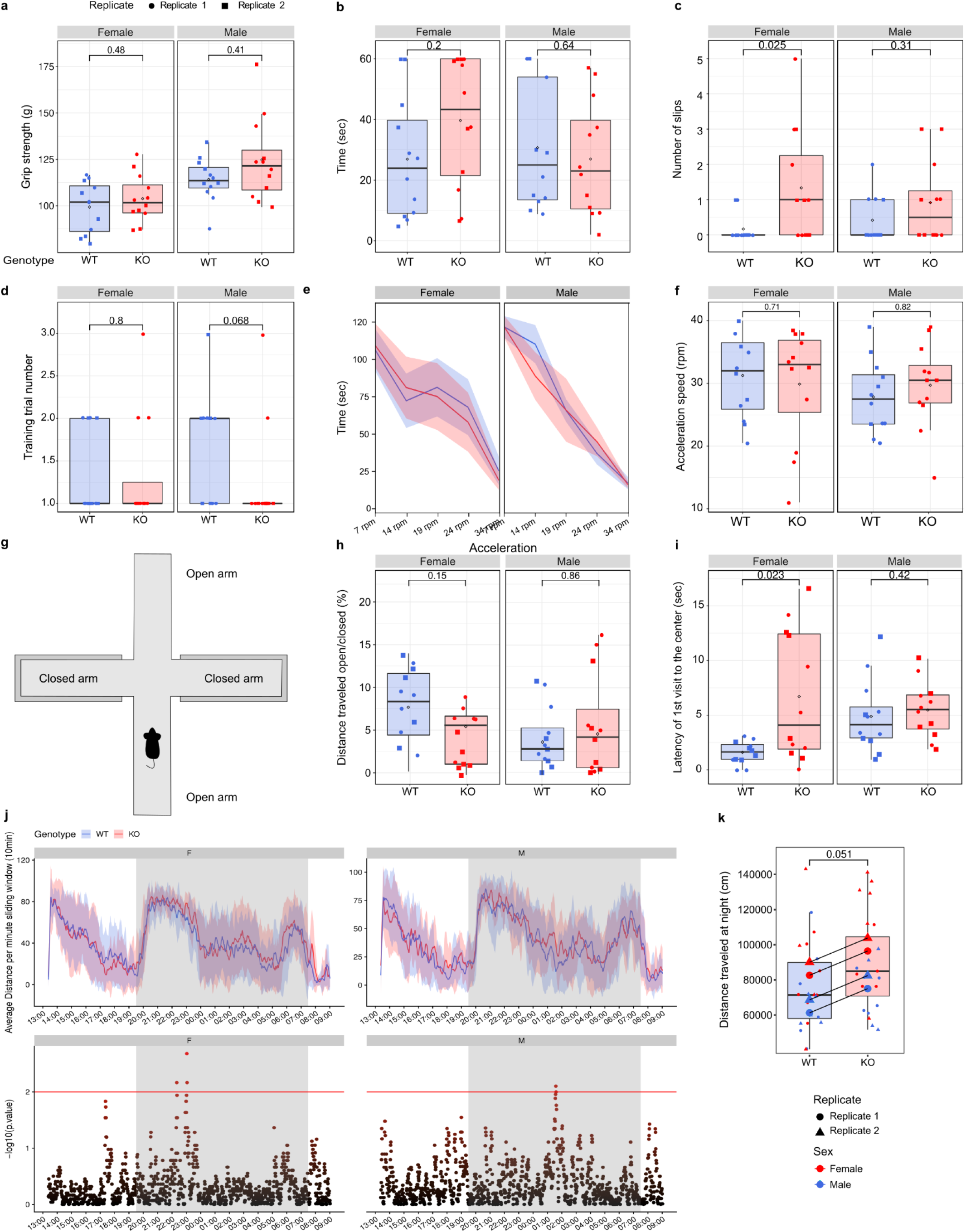
Activity, motor coordination and balance tests in adult mice. (a) Distributions of maximal peak force developed as a measure of adult forelimb grip strength. Each value corresponds to the average of three replicate measures per mouse. (b,c) Time needed to perform the beam balance task (b) and number of slips from the beam per animal (c). (d-f) Rotarod performance analysis with training trial number needed to perform the task (d), time-based performance during the constant speed sessions (e), and performance during the acceleration session (f). (g) Schematic representation of the elevated plus maze experiment. Red rectangles describe the open-arm ends. (h) Distributions of the ratio of the time spent in the open vs. closed arms of the plus maze. P-values from Wilcoxon rank-sum tests. (i) Latency for the entrance to the central zone of the plus-maze. (j) Spontaneous locomotor activity of WT and KO animals during a continuous 23 h period. Top: Average distance traveled per genotype in 10 min (sliding window of 1 min). Bottom: log10 p-value for each 10-min interval, calculated through permutation tests. The horizontal red line describes the significance threshold, and the gray area marks the light-off/dark phase of the night cycle. The X-axis describes time and corresponds to the 24-hour notation as hh:mm. (k) Distance traveled for WT and KO mice during the 12-hour night period (lights off). P-values from Wilcoxon rank-sum tests (a-i) and two-way ANOVA tests with replicate and genotype as factors (k).

**Supplementary Fig 8.**
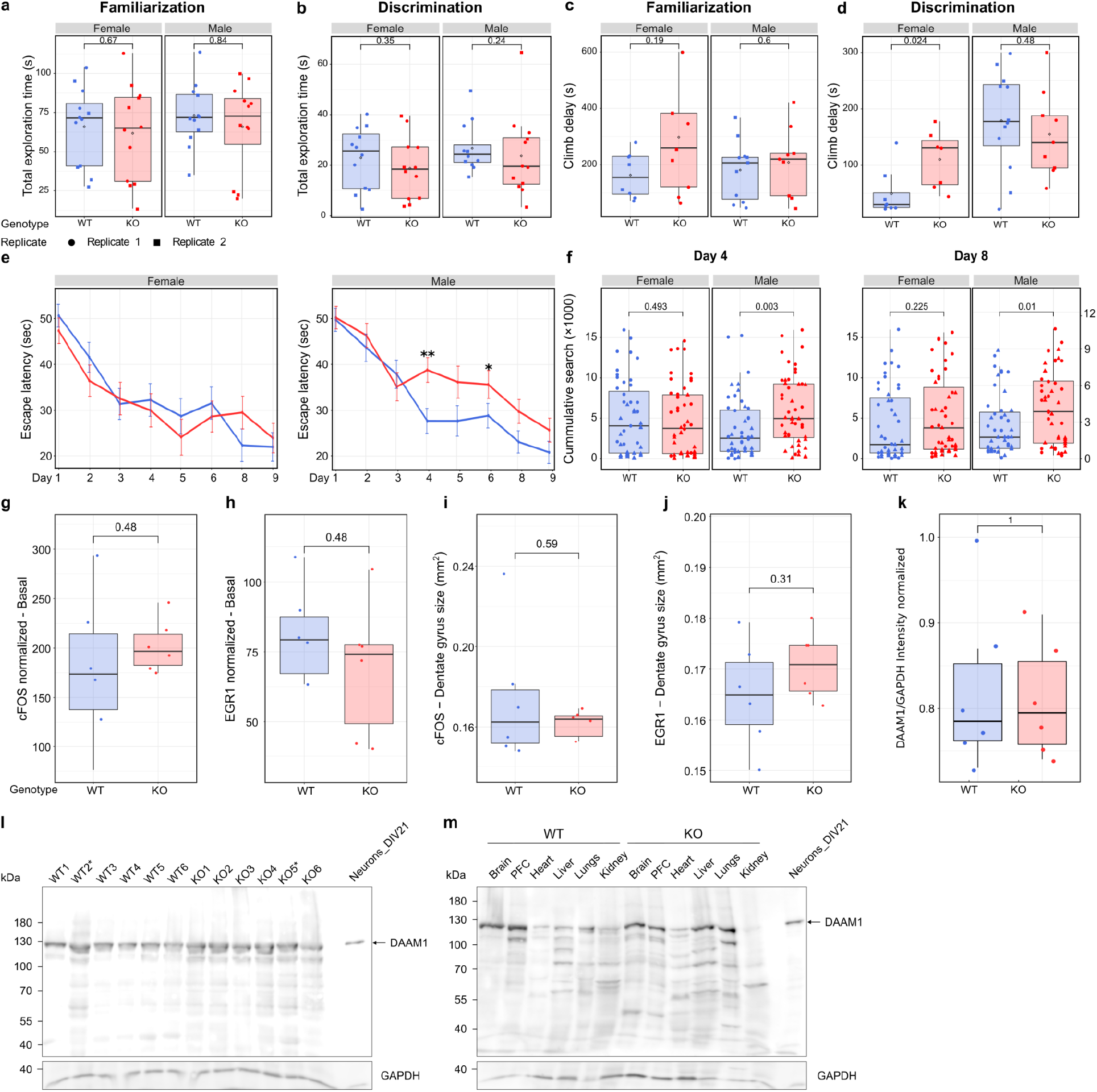
Cognition and memory-related tests. (a,b) Total exploration time is defined as object sniffing during the familiarization (a) or discrimination (b) phases of the NOR experiment performed. (c,d) Object climb delay during the NOR familiarization (c) or discrimination phase (d). (e) Latency to find the escape route (hidden platform) in seconds during the acquisition phase for females (upper panel) and males (lower panel). (f) Quantification of the cumulative index during the 4th and 8th days of the acquisition phase. Dots represent the performance of one mouse during one trial. (g-j) Quantification of cFOS (g) and EGR1 (h) positive nuclei in the hippocampal dentate gyrus at basal conditions. Number of IEG-positive nuclei was normalized to the dentate gyrus size ([i] and [j], respectively). One dot represents an average of 3-6 coronal views of the hippocampus analyzed per animal, performed in males. (k,l) Western blot (k) and associated quantification (l) of DAAM1 protein expression in cerebellum and motor cortex. The asterisk indicates samples derived from male animals. The relative intensity of DAAM1 was normalized to GAPDH. Dots represent values measured on individual cerebellum. (m) Western blot of DAAM1 protein isoforms in multiple tissues. P-values from Wilcoxon rank-sum tests (a-d,h-o,q) and two-way ANOVA tests with replicate and genotype as factors (e,f).

**Supplementary Figure 9.**
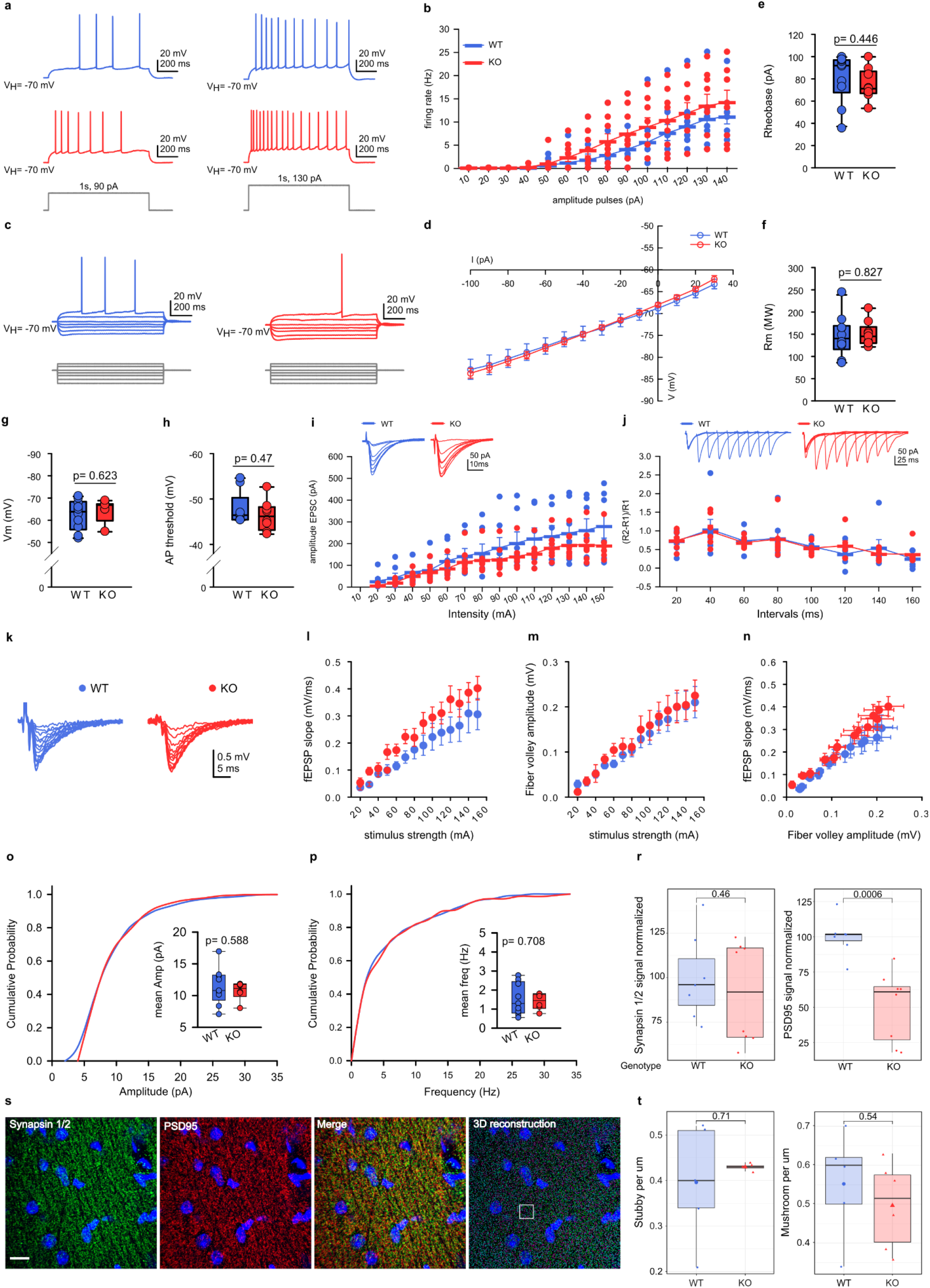
Intrinsic membrane properties and synaptic plasticity study in CA1 pyramidal cells. (a) Representative traces of action potential firing frequency induced by depolarizing current steps at 90 pA (left) and 130 pA (right) (1s duration) in WT (blue traces) and KO (red traces) cells. (b) Average firing frequency versus injected current relationship for representative WT and KO cells. No statistically significant differences were found by two-way repeated measures ANOVA tests (WT: n= 10, N= 4; KO: n= 9, N= 4). Error bars indicate SEM. (c) Voltage responses of representative WT (blue traces) and KO (red traces) cells to hyperpolarizing and depolarizing current steps. (d) V/I curves of WT (blue circles) and KO (red circles) cells. Voltage (V) measures in response to hyperpolarizing and depolarizing current (I) steps (WT: n= 10, N= 4; KO: n= 9, N= 4). Error bars indicate SEM. (e-h) Box plots of rheobase (Rh, [e]), membrane resistance (Rm, [f]), resting membrane potential (Vm, [g]), and threshold action potential (th AP, [h]). P-values from Welch’s t-test (e, f) and Mann-Whitney (g, h). (WT: n= 10, N= 4; KO: n= 9, N= 4). (i-j) Summary of the EPSC experiment. (i) Example evoked EPSC traces by stimulation of the SC pathway applying different intensities (pA) in WT (top, blue traces) and KO (bottom, red traces) CA1 cells (upper panel). EPSC amplitude values for a given range of stimulus intensities in WT (n = 7, N= 3) and KO (n = 8, N= 3) cells (lower panel). No statistically significant differences were found by two-way repeated measures ANOVA tests. Error bars indicate SEM. (j) Representative pair of evoked EPSC traces for different intervals (ms) of stimulation of the SC axons in WT (top, blue traces) and KO (bottom, red traces) CA1 cells (upper panel). Paired-pulse ratios calculated from EPSC amplitudes ((R2-R1)/R1, where R1 corresponds with the first response and R2 second response) across varying inter-stimulus intervals showed unchanged facilitation between WT (n = 7, N= 3) and KO (n = 8, N= 3) cells (lower panel). No statistically significant differences were found by two-way repeated measures ANOVA. Error bars indicate SEM. (k, l, m, n) KO mice do not modify the basal synaptic transmission at CA1 synapses. (k) Representative fEPSPs recorded in the stratum radiatum and evoked by stimulation of the SC pathway with different intensities in control (blue circles) and in the KO response (red circles). (l) fEPSP slopes were not statistically significant between control (blue circles, n= 6, N= 3) and KO responses (red circle; n= 6, N= 4) for a given range of stimulus intensities. (m) Fiber volley amplitudes are similar between experimental conditions (control, blue circle, n= 6, N= 3; KO, red circle, n= 5, N= 4) for a given range of stimulus intensities. (n) Input/output relationships for control (blue circles) and KO responses (red circles). Statistical analysis of the basal synaptic transmission with two-way RM ANOVA shows no significant interaction. (o,p) Cumulative probability plot and box plot of amplitude (o) and frequency (p) isolated miniature EPSC events recorded in pyramidal cells in WT (n= 9, N= 3, Events= 1703) and KO (n= 6, N= 3, Events= 1040) CA1 cells. (r) Distributions of the percentage of Synapsin 1/2 and PSD95 signal normalized to the control mean. P-values from Wilcoxon rank-sum tests. (s) Representative images of Synapsin 1/2 (presynaptic), PSD95 (postsynaptic) markers, and 3D Imaris reconstruction of the merged images from the CA1 hippocampal region of a PD22 control mouse. (t) Number of stubby (left) and mushroom (right) spines per 1 μm of neurite from biocytin-stained CA1 pyramidal neurons of the hippocampus in WT and KO mice. N = 6 mice for each genotype, 100 μm of dendrites per cell. One dot represents one averaged value per mouse. Lines represent the relation between animals analyzed in experimental replicates. P-values from Wilcoxon rank-sum tests.

**Supplementary Figure 10.**
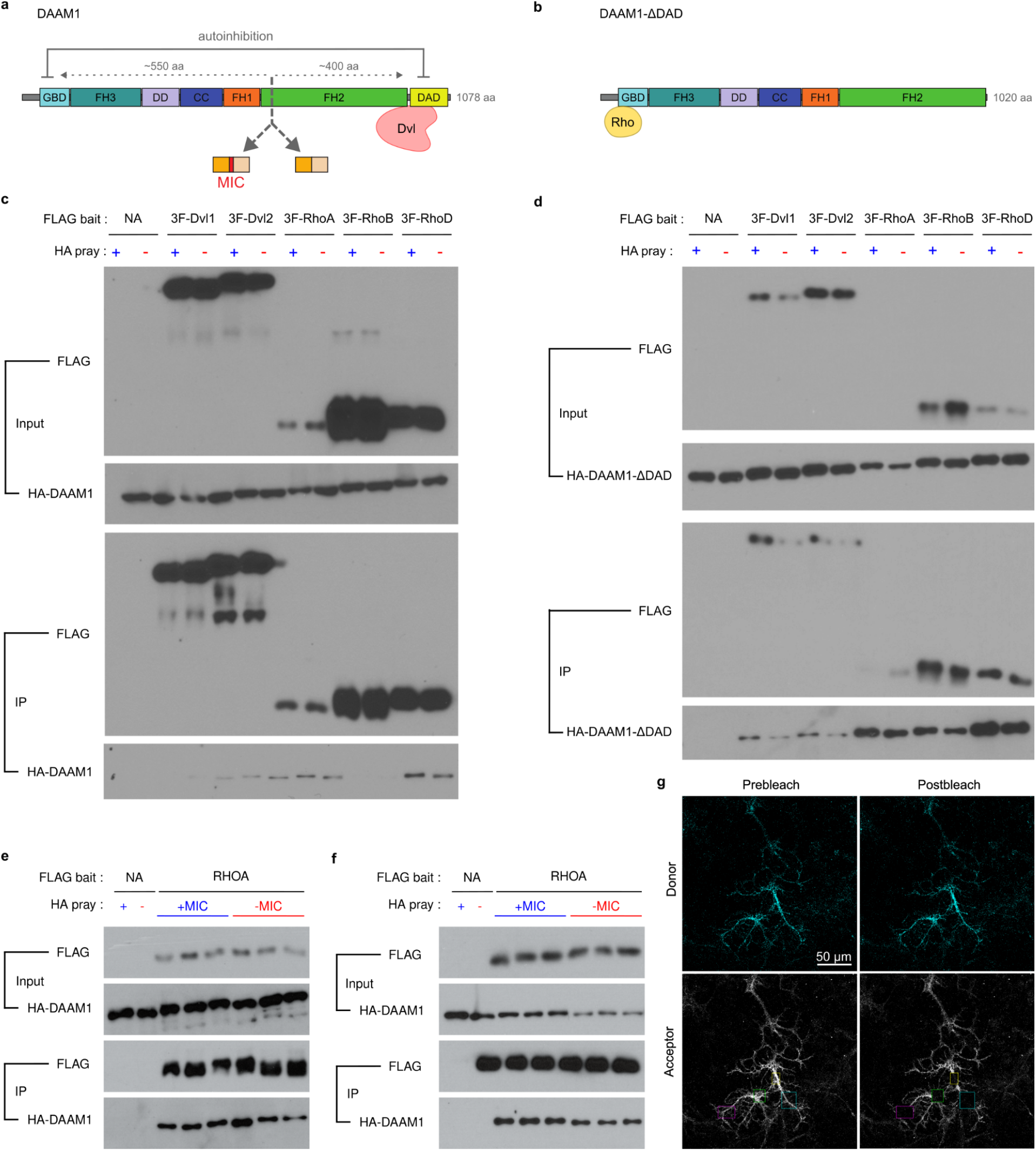
Daam1-MIC removal impairs the RHOA/ROCK signaling pathway. (a) Schematic representation of the DAAM1 protein together with its domains and known interactors. The predicted distance from the microexon inclusion region to the RHOA interacting domain (GBD) and the DVL interacting domain (DAD) is in gray. (b) DAAM1-ΔDAD corresponds to the DAD truncated variant that releases the autoinhibition allowing RHOA interaction. Domain nomenclature as in Figure 1e. (c-f) 293T cells were transfected with HA-tagged DAAM1 (c) or DAAM1-ΔDAD (d-f) constructs, with or without the microexon, together with the 3xFlag-tagged protein indicated. Immunoprecipitation was performed with anti-Flag antibody. (g) Representative images of RhoA2G acceptor photobleaching in differentiated neuronal cells (DIV21). Colored boxes represent photobleached regions of interest.

**Supplementary Figure 11.**
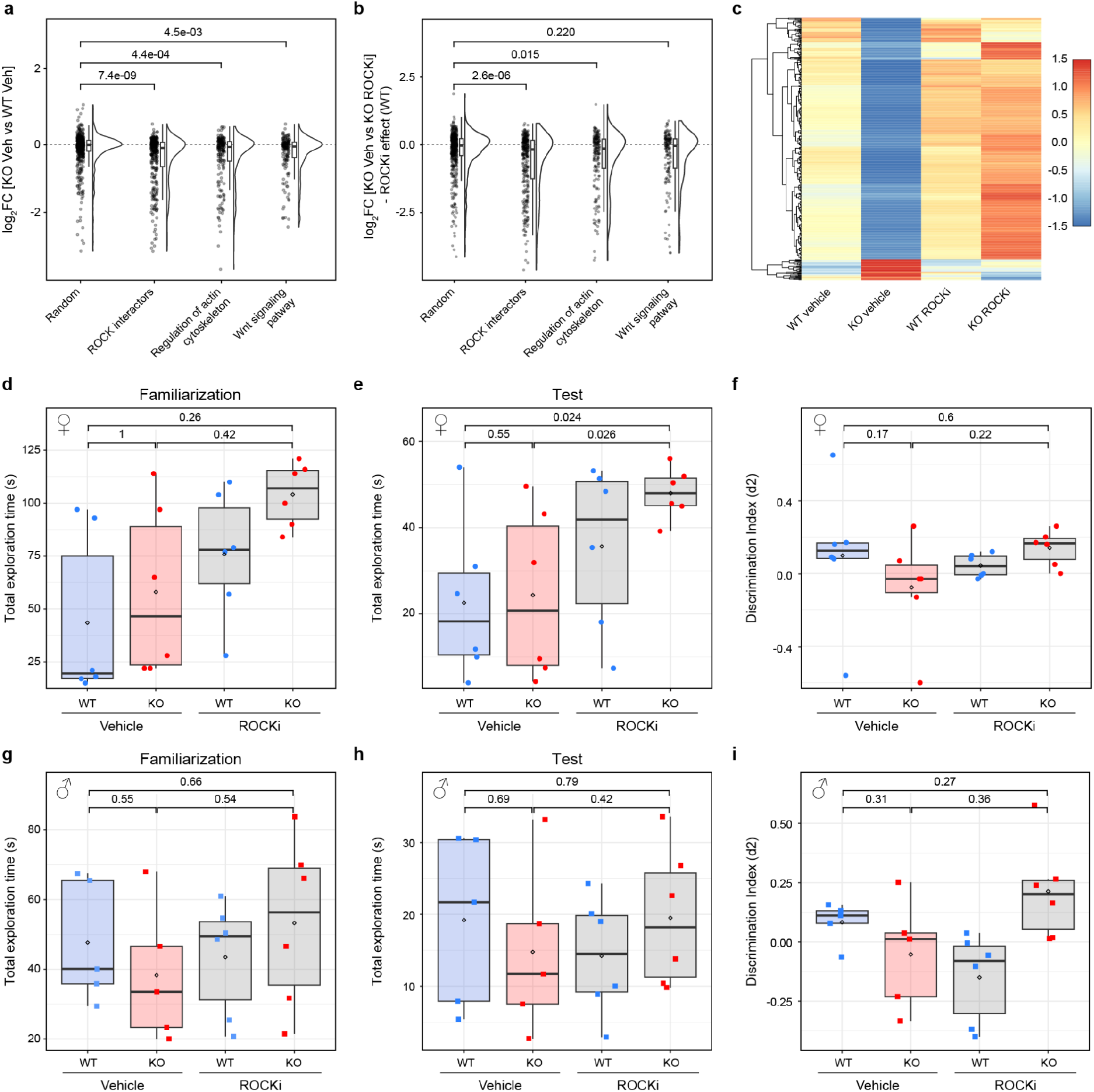
Rescue upon ROCK pathway inhibition. (a,b) Distribution of gene expression changes (log2 fold change) between Daam1-MIC KO and WT cells under control conditions for gene sets of interest: a random set of genes, proteins that interact with DAAM1 based on the STRING database, and two KEGG pathways where DAAM1 is involved (regulation of actin cytoskeleton and Wnt signaling pathway). (c) Relative expression of genes with a rescue pattern across the four experimental conditions. Each row corresponds to a rescued gene, and the values correspond to the z-score of the average of the three replicates per condition. (d-i) Summary of the NOR experiment performed after the ROCK inhibitor treatment, where the total exploration time is defined as object sniffing during the familiarization (a and d) or discrimination (b and e) phases, and (c and f) distribution of discrimination index (d2) values quantified during the NOR discrimination phase. The discrimination index corresponds to: (Novel Object Exploration Time - Familiar Object Exploration Time)/ Total Exploration Time. The upper panel (a-c) describes the performance for females, and the bottom panel (d-f) for males is indicated on the left side. One dot describes the performance of one animal. P-values from Wilcoxon rank-sum tests.

**Supplementary Figure 12.**
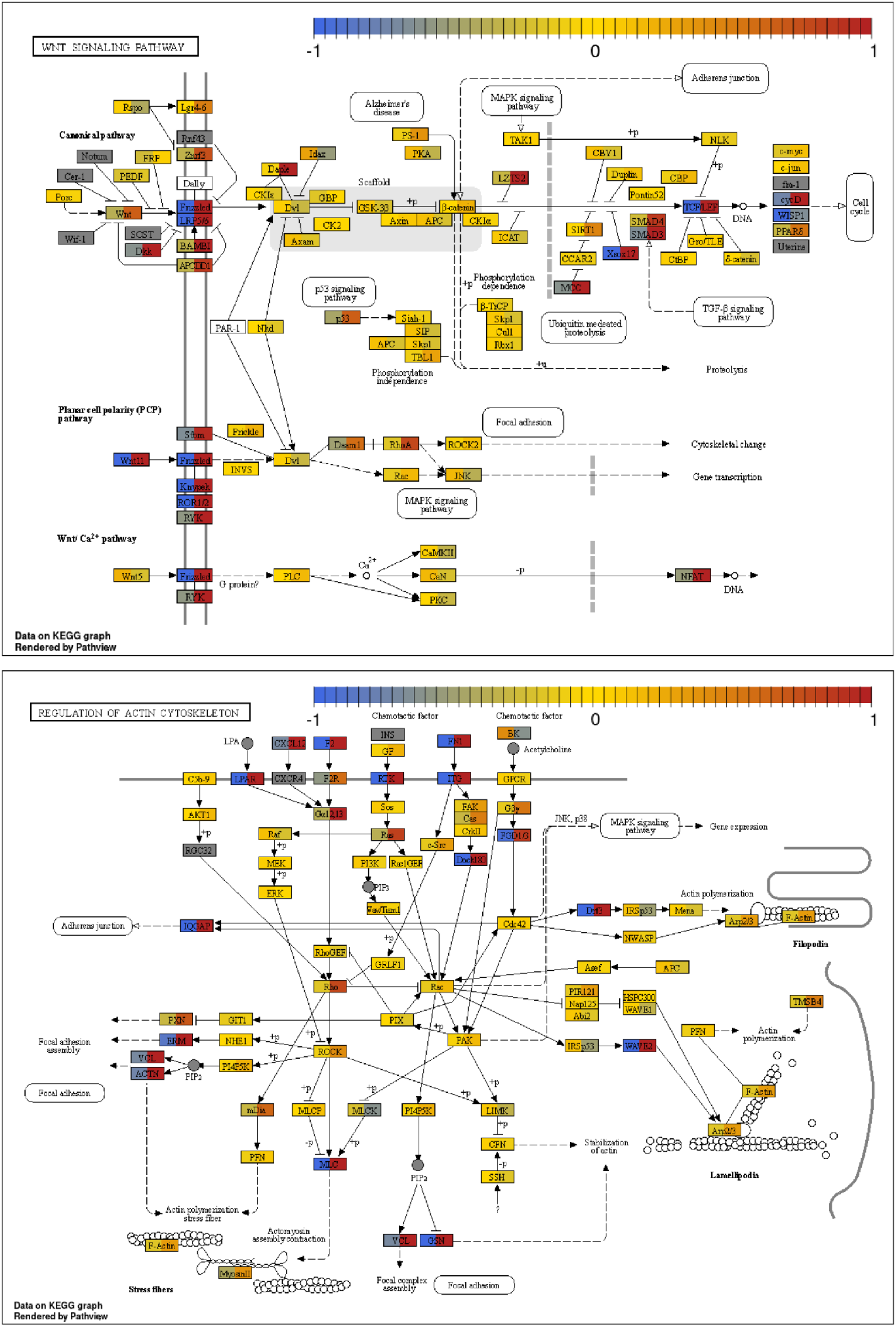
RNA-seq rescue of selected KEGG pathways upon ROCK inhibition. KEGG pathway for the Wnt signaling pathway (upper panel), and the regulation of actin cytoskeleton (lower panel) where elements are colored based on log2 fold change between control WT and KO neurons (left side), and between KO neurons treated with Vehicle and ROCK inhibitor Y-27632 (right side).

## Supplementary Tables

**Supplementary Table S1.**
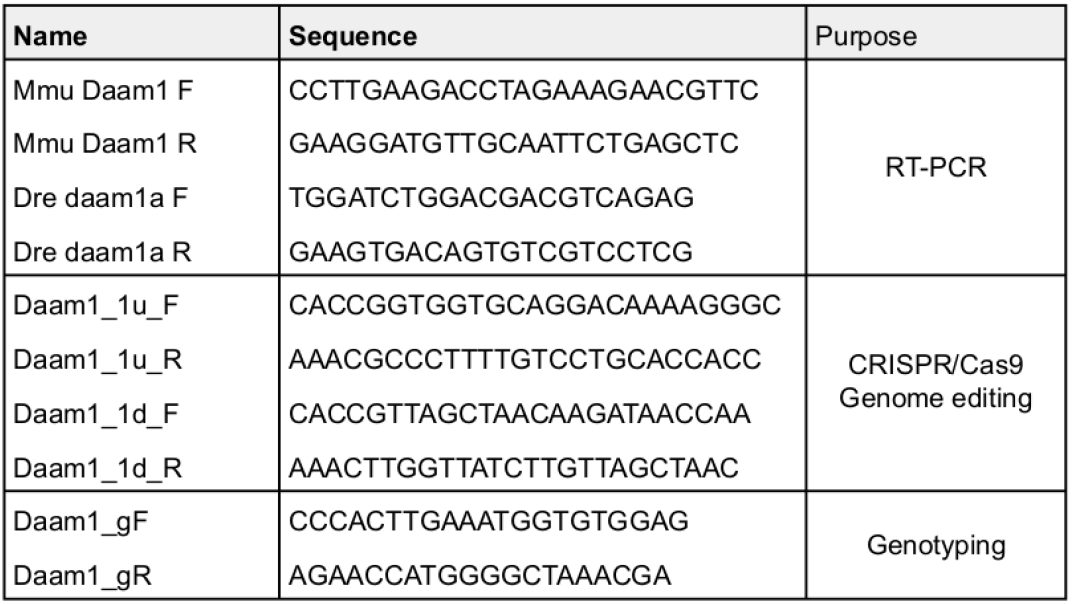

**Supplementary Table S2.**
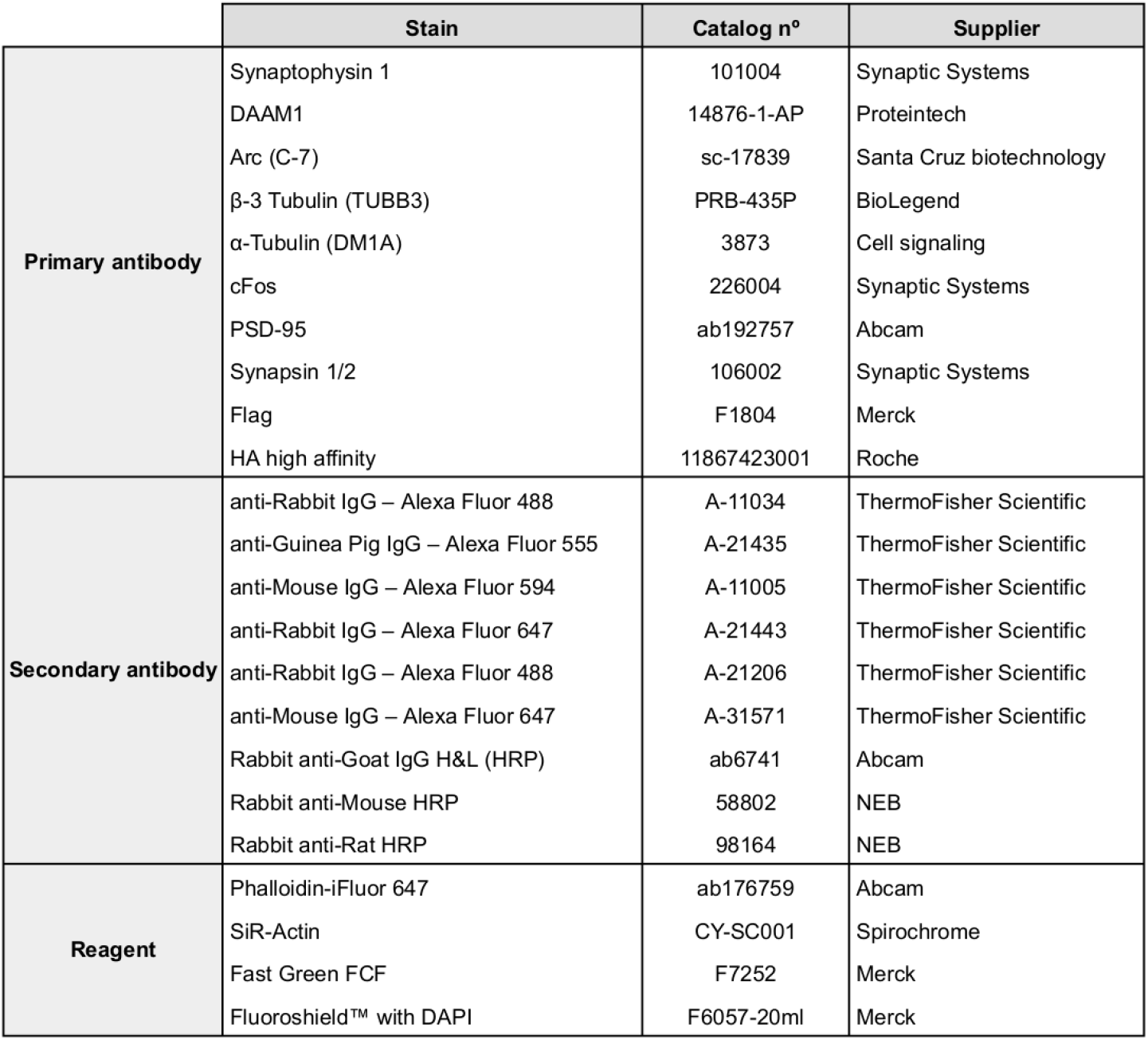

|  | Stain | Catalog n° | Supplier |
| --- | --- | --- | --- |
| **Primary antibody** | Synaptophysin 1 | 101004 | Synaptic Systems |
|  | DAAM1 | 14876-1-AP | Proteintech |
|  | Arc (C-7) | sc-17839 | Santa Cruz biotechnology |
|  | β-3 Tubulin (TUBB3) | PRB-435P | BioLegend |
|  | α-Tubulin (DM1A) | 3873 | Cell signaling |
|  | cFos | 226004 | Synaptic Systems |
|  | PSD-95 | ab192757 | Abcam |
|  | Synapsin 1/2 | 106002 | Synaptic Systems |
|  | Flag | F1804 | Merck |
|  | HA high affinity | 11867423001 | Roche |
| **Secondary antibody** | anti-Rabbit IgG – Alexa Fluor 488 | A-11034 | ThermoFisher Scientific |
|  | anti-Guinea Pig IgG – Alexa Fluor 555 | A-21435 | ThermoFisher Scientific |
|  | anti-Mouse IgG – Alexa Fluor 594 | A-11005 | ThermoFisher Scientific |
|  | anti-Rabbit IgG – Alexa Fluor 647 | A-21443 | ThermoFisher Scientific |
|  | anti-Rabbit IgG – Alexa Fluor 488 | A-21206 | ThermoFisher Scientific |
|  | anti-Mouse IgG – Alexa Fluor 647 | A-31571 | ThermoFisher Scientific |
|  | Rabbit anti-Goat IgG H&L (HRP) | ab6741 | Abcam |
|  | Rabbit anti-Mouse HRP | 58802 | NEB |
|  | Rabbit anti-Rat HRP | 98164 | NEB |
| **Reagent** | Phalloidin-iFluor 647 | ab176759 | Abcam |
|  | SiR-Actin | CY-SC001 | Spirochrome |
|  | Fast Green FCF | F7252 | Merck |
|  | Fluoroshield™ with DAPI | F6057-20ml | Merck |

## Supplementary Methods

### Mouse embryonic stem cell culture

mESC were grown on 0.1% gelatine-coated (Millipore, ES-006-B) plates (Thermo Scientific Nunc Cell-Culture Treated Multidishes, 140675) with mESC media containing 10% fetal bovine serum (FBS) and leukaemia inhibitory factor (LIF). Gelatine coating was performed for a minimum of 5 minutes before plating the cells. mESC media was generated by the CRG Tissue Engineering Unit and consisted of: Glasgow’s Minimum Essential Medium (GMEM) BHK-21 (Gibco, 21710-025) supplemented with 10% fetal bovine serum (Seralab, A1060013 EU-000-H), Minimum Essential Medium non-essential amino acids solution (Gibco,11140-050), 1mM L-Glutamine (Gibco, 25030-024), 0.5mM Sodium Pyruvate (Gibco, 11360-070), 0.1mM 2-Mercaptoethanol (Millipore, ES-007-E) and recombinant mouse LIF protein 1000U/ml ESGRO (Millipore, ESG1107). mESCs were routinely passaged using 1x TrypLE Express Enzyme (Gibco, 12605028).

### Neuronal differentiation

We followed the protocol reported by Bibel et al., 2007 with slight modifications. In brief, mESCs were harvested by trypsinization with TrypLE Express for 5 min at 37°C (HeracellTM 240i CO2 Incubator). Trypsin was quenched with an equal volume of Embryoid Body (EB) medium (10% FBS, 1% Non-Essential Amino Acids, 1% Penicillin and Streptomycin, 1% GlutaMax, 1% Sodium Pyruvate, 0.1% B-mercaptoethanol, 86% DMEM High Glucose) and cells were counted manually using a haemocytometer chamber. mESCs were plated at a density of 4x106 cells per low attachment bacteriological petri dish (10 cm Ø) in 15 ml of EB medium. This marks the start of the experiment and is further referred to as Day In Vitro -8 (DIV-8). Cells were cultured at 37°C with 5% CO2. On day 2 medium with the Embryoid Bodies (EB) was transferred to the falcon tube, and after 5 min supernatant was aspirated and pelleted EBs were resuspended in fresh EB medium (DIV-6). Resuspended EBs were dispensed into new bacteriological Petri dishes (10 cm diameter) in EB medium volume up to 15 ml per dish, and incubated as earlier. On day 4 (DIV-4) and day 6 (DIV-2) the medium was changed as before but using EB medium supplemented with 5 μM retinoic acid. On day 8 (DIV0) EBs were collected as before and washed twice with 10 ml of PBS. Subsequently, EBs were resuspended in 1 ml of medium (0.05% Trypsin, 0.05% EDTA dissolved in PBS) and incubated with constant shaking for 3 min in a 37°C water bath. Trypsinized EB’s were quenched with 1 ml of EB medium and cells were pelleted down using Eppendorf 5810R Centrifuge (180 g for 5 min). The supernatant was aspirated and the cells were resuspended in 5 ml of N2 medium and filtered through a 40μm cell strainer. The cells were counted as before and plated at the density of 1.5x105 per 13 mm glass coverslip (VWR 631-1578) placed in 24 well plates (Sigma Aldrich, 11243217001) coated with poly-D-Lysine followed by laminin (Roche). Cells were cultured as described above in the incubator set up at 37°C with 5% of CO2. After 2, 24 and 48 h from plating, the N2 medium was changed. Consecutively, media was changed to B27 after 72 h and fresh B27 media was provided every second day. Neuronal differentiation was conducted up to DIV23.

### Behavioral and locomotor tests in a neonatal mice

A battery of behavioral and motor tests to probe early post-natal neurodevelopment was performed as described in Feather-Schussler and Ferguson (2016) and Roper et al. 2021, with some adjustments. In particular, we performed tThe following tests were performed at PDs 4, 7, 10 and 14, unless stated otherwise:

#### Pivoting and walking

Pivoting is a voluntary exploratory behavior displayed by young mice before the forward locomotion. To measure pivoting, mice were placed on a flat surface and allowed to move freely for 1 min. The number of times the animals made a 90° turn was recorded. Pivoting was measured based on the body axis with the help of a cross marking 90° angles on the experimental surface. The latency to walk in a straight line after the end of the pivoting behavior was also measured. Two trials were performed per animal.

#### Righting reflex

Mice (PD4 or PD7) were placed on their back on a padded table top. The time taken for the animals to right themselves back to four paws through 180° was measured for a maximum of 1 min. The direction of turning was also recorded. The experiment was performed three times per animal and the mean calculated.

#### Preyer’s reflex

The Preyer’s reflex is a startle response triggered by sharp auditory stimuli and is used to assess hearing in rodents (Jero, Donald E. Coling, Anil K. Lal 2001). The experimenter made a sharp clapping sound by stretching and releasing a rubber glove onto the hand in the proximity of the mouse and recorded the presence of rapid whole-body movement.

#### Front-limb suspension

To measure forelimb strength, mice were allowed to grasp with both forepaws on a horizontal bar suspended above a padded drop zone. The latency to fall was measured in three trials per animal and the mean was calculated.

#### Hindlimb suspension

To measure hindlimb strength, mice were lowered into a 50 ml conical tube and released with their hindlimbs hung over the rim. The latency to fall was measured for up to 30 s. Hindlimb posture upon falling was scored 0-4 based on the limb spread, as described in Feather-Schussler and Ferguson (2016). The experiment was performed only once unless the animal fell down immediately due to bad placement.

#### Grasping reflex

The animals were held by the scruff of the neck and each paw was touched by a toothpick to elicit the grasping reflex. Performance was scored 0-4, assigning one point per paw in the presence of grasping. Left and right paw preference was also noted.

#### Cliff aversion

Cliff aversion tests the labyrinth reflexes, as well as normal strength and coordination. Mice were placed on top of a box elevated ∼10 cm above the surface, with their snout and part of their forepaws just over the edge. The presence of aversive movement away from the cliff within the subsequent 30 s was recorded. If the pup fell down, one additional trial was performed.

#### Negative geotaxis

Negative geotaxis is an automatic vestibular response to geogravitational stimuli and is used to measure motor coordination in pups. Mice were placed head-facing uphill on a plastic platform with an inclination of 45° covered with Surface Protector Paper and spunlace wipes (VWR), except for PD 4 mice, for which the incline was adapted to 30°. After ∼5 s, the pups were turned by 180° to face downhill, and their movement was observed for 1 min. Two trials were performed, turning the animal in opposite directions to the start position to avoid left-right bias. Animals were given 0.5 points per 45° of turning. If a full 180° turn was performed, the latency was also recorded. Left and right turning preferences were also noted.

#### Homing test

Homing was performed as described in Roper et al. 2021 at PD14 with adjustments. A pup is removed from the home cage and is placed in the corner of a new clean cage (12.5 × 45 cm, w × h) facing the wall. The new cage is filled with clean wood shavings, and familiar nesting material from its home cage is provided in the opposite corner. The time for the pup to reach the area containing the nesting material is recorded in seconds, with a maximum time of 2 min.

### Spontaneous basal locomotor activity in adult mice

Spontaneous basal locomotor activity during day and night in an open cage was measured for 23 h using an infrared Actimeter (PANLAB SA, Spain). Individual mice were placed in the open field (25 × 25 cm), and their position in time was recorded based on the disruption of infrared beams in the x and y axes. The arena was covered with a layer of wood shavings, and sufficient water and food was provided.

### Novel Object Recognition (NOR) test

The NOR test measures recognition memory based on the visual paired-comparison paradigm (Leger et al. 2013; Lueptow 2017). The experiment consists of three sessions carried out on consecutive days: habituation, familiarisation, and discrimination. The experiment was performed in a 38.5 cm × 38.5 cm arena with 38.5 cm high dark plastic walls and an open top. The arena was illuminated from the top, and the animals were tracked using video recording and the Smart 3.0 software (System Motor Activity Record and Tracking, PANLAB SA, Spain). To minimize stressors interfering with the experiment, a curtain was mounted separating the arena from the experimenters. The arena and the objects were cleaned with 70% ethanol between each mouse to remove odors. Cages were transferred to the experimental room 30 min before the trials to acclimatize. The test was carried out under slightly aversive conditions (white light 50 lux).

#### Habituation

On the first day, animals were placed into the arena facing the wall and were recorded for 5 min. Their movement was measured separately by the Smart 3.0 software in a central (20 × 20 cm) and a peripheral zone.

#### Familiarization

On the second day, two identical objects (A and B) were placed in the centre of the box, 18 cm apart. Animals were placed into the box facing the wall and were recorded for 10 min. The time spent exploring each object was measured manually using a timer. The latency to climb on top of the objects was also noted. The exploration was defined as the time spent sniffing the object from a close distance, excluding the time spent climbing on the object. Exploration threshold was established at 20 sec.

#### Discrimination

On the third day, one of the objects was replaced with a novel object of a different shapes and colours. Animals were placed into the box facing the wall as before and were recorded for 5 min. The time spent exploring each object was measured manually using a timer. The latency to climb on top of the old and novel objects was noted. Discrimination and preference indices were calculated for familiarisation and discrimination for each animal based on the manual exploration time recordings, as described in Lueptow et al. (2017).

### Elevated Plus Maze test

Anxiety-related behavior was measured in the Elevated plus maze, as described by (Walf and Frye 2007), with some modifications. The apparatus consisted of a cross-shaped platform with four arms (30 cm × 5 cm), elevated 40 cm off the ground. Two arms were open and two were enclosed by 15 cm high black methacrylate walls. The open arms create an aversive environment for the mouse, while the dark enclosed space of the closed arms is considered safe. The number of open arm entries and the relative time spent in the open arms is indicative of anxiety-related behavior. Animals that avoid open arms are considered more anxious. Mice were lowered onto the end of the same closed-arm facing the walls and their movement was recorded for 5 min using the Smart 3.0 software. Separate zones were set up for each arm, the ends of the arms (5 cm × 5 cm) and the center of the cross (5 cm × 5 cm). Measurements included the percentage of the time, entries, distance traveled, and average speed in each zone. Rearings and head dippings from the central zone were recorded manually. The arena was cleaned with 70% ethanol between each animal’s experiment to remove odors.

### Morris Water Maze test

Learning and visual-spatial memory were tested in the Morris water maze (Morris 1984), with some modifications (Vorhees and Williams 2006). In this experiment, mice must learn the spatial location of a submerged platform to escape from the water by building a cognitive allocentric map with the help of external visual cues. A metal tank (150 cm in diameter) was filled with water (22-23°C) with added white, non-toxic finger paint to conceal the location of the platform. The platform (12 cm in diameter) was submerged 0.7 cm below the water surface. Four quadrants (NE, NW, SE, SW) and a platform zone were defined using the Smart 3.0 software, with the platform located in the center of one quadrant (NE). A curtain was mounted around the tank to separate it from the experimenters and minimize external visual cues. Three distal visual cues, a square, a triangle and a circle were placed on the curtains around the tank at equal distances, ∼30 cm above the water surface. The experiment consists of the following phases: training trials, removal, cued trials, and reversal training.

#### Training trials

During training, the mice learn the task and the location of the platform. Four 1-min training trials were performed each day, with an inter-trial interval of ∼1 h. Nine training days were performed to achieve sufficient learning. Mice were released into the water. The latency to reach the platform was recorded and the animal was removed from the water. 5 to 15 s platform localization learning and positive reinforcement was used by placing the animal on the platform depending on performance (goal accomplishment or lack of it).

#### Removal

the platform is removed from the tank and the animals are tested for 1 min 24 h after the last training. Animals were released from the location furthest away from the original platform (SW). Mice that remember the location of the platform are expected to spend more time in the platform quadrant (NE).

#### Cued trials

In these guided learning trials, the platform is placed back to its original location and is marked by a local visual cue. Animals have an inherent tendency to swim towards the flag. Therefore, these trials can detect issues with swimming, visual perception and/or motivation to perform the task. A flag (∼10 cm above the water surface) fixed to a metal bar was used as a local cue. Distant visual cues were removed. Two 1-min cued trials were performed ∼1 h after the removal session.

#### Reversal training

During a spatial reversal, the platform is placed opposite to its original location (SW). Mice must flexibly re-learn the spatial location of the platform. Four trials were performed for 2 days, as described in the training trials.

In all trials, the latency to reach the platform and platform crossovers were measured. Animals were tracked using the Smart 3.0 software, allowing measurement of time, speed and distance traveled in each quadrant. Distances from the platform were used to calculate cumulative search error and mean proximity as additional measurements of spatial learning (Pereira & Burwell, 2015). Floating was also measured manually.

### Grip strength test

The grip strength experiment is a measurement of neuromuscular function and limb muscle strength. Mice were held by the tail and lowered onto the metal grid of the apparatus (Grip Strength Meter, Bioseb, France), allowing them to grip with their paws. Mice were next pulled backwards along the grid at a consistent speed, while the apparatus measured the force exerted by the animals on the grid. Grip strength was measured for forelimbs through 3 consecutive trials. Best, average, and mean grip strengths were calculated.

### Rotarod test

To evaluate coordination and balance, we employed the Rotarod apparatus (PanLab Rotarod LE8200, Spain) based on (Deacon 2013). First, animals were trained to walk on the rod at 4 rpm. Training sessions were performed until the mouse managed to walk on the rod for 1 min. After training, test sessions were performed at five constant speed settings (7, 14, 19, 24, 34 rpm). Two trials were carried out per speed, with a maximum length of 2 min. Finally, two trials were performed where the rotation speed accelerated constantly from 4 to 40 rpm in 60 s.

### Beam Balance test

Balance, coordination, and vestibular function were measured in the Beam Balance experiment (Luong et al. 2011). Mice were placed in a standing position in the center of a narrow wooden beam (1 cm × 50 cm) elevated 45 cm above the ground. They were scored 0-3 based on how far they walked on the beam within 1 min (0: falls off; 1: <10 cm; 2: >10 cm; 3: reaches the end). Falls and the number of slips were also recorded, as well as the latency to reach one of the ends and the direction of movement (left/right).

## References

Barbosa-Morais, N. L., M. Irimia, Q. Pan, H. Y. Xiong, S. Gueroussov, L. J. Lee, V. Slobodeniuc, et al. 2012. “The Evolutionary Landscape of Alternative Splicing in Vertebrate Species.” Science 338 (6114): 1587–93. 10.1126/science.1230612.

Bibel, Miriam, Jens Richter, Emmanuel Lacroix, and Yves-Alain Barde. 2007. “Generation of a Defined and Uniform Population of CNS Progenitors and Neurons from Mouse Embryonic Stem Cells.” Nature Protocols 2 (5): 1034–43. 10.1038/nprot.2007.147.

Bibel, Miriam, Jens Richter, Katrin Schrenk, Kerry Lee Tucker, Volker Staiger, Martin Korte, Magdalena Goetz, and Yves-Alain Barde. 2004. “Differentiation of Mouse Embryonic Stem Cells into a Defined Neuronal Lineage.” Nature Neuroscience 7 (9): 1003–9. 10.1038/nn1301.

Bramham, Clive R., Maria N. Alme, Margarethe Bittins, Sjoukje D. Kuipers, Rajeevkumar R. Nair, Balagopal Pai, Debabrata Panja, et al. 2010. “The Arc of Synaptic Memory.” Experimental Brain Research 200 (2): 125–40. 10.1007/s00221-009-1959-2.

Calarco, John A., Simone Superina, Dave O’Hanlon, Mathieu Gabut, Bushra Raj, Qun Pan, Ursula Skalska, et al. 2009. “Regulation of Vertebrate Nervous System Alternative Splicing and Development by an SR-Related Protein.” Cell 138 (5): 898–910. 10.1016/j.cell.2009.06.012.

Ciampi, Ludovica, Federica Mantica, Laura López-Blanch, Jon Permanyer, Cristina Rodriguez-Marín, Jingjing Zang, Damiano Cianferoni, et al. 2022. “Specialization of the Photoreceptor Transcriptome by Srrm3-Dependent Microexons Is Required for Outer Segment Maintenance and Vision.” Proceedings of the National Academy of Sciences 119 (29): e2117090119. 10.1073/pnas.2117090119.

Cingolani, Lorenzo A., and Yukiko Goda. 2008. “Actin in Action: The Interplay between the Actin Cytoskeleton and Synaptic Efficacy.” Nature Reviews Neuroscience 9 (5): 344–56. 10.1038/nrn2373.

Consolati, Tanja, Gil Henkin, Johanna Roostalu, and Thomas Surrey. 2022. “Real-Time Imaging of Single ΓTuRC-Mediated Microtubule Nucleation Events In Vitro by TIRF Microscopy.” In Microtubules, edited by Hiroshi Inaba, 2430:315–36. Methods in Molecular Biology. New York, NY: Springer US. 10.1007/978-1-0716-1983-4_21.

Dillon, Christian, and Yukiko Goda. 2005. “THE ACTIN CYTOSKELETON: Integrating Form and Function at the Synapse.” Annual Review of Neuroscience 28 (1): 25–55. 10.1146/annurev.neuro.28.061604.135757.

Doench, John G, Nicolo Fusi, Meagan Sullender, Mudra Hegde, Emma W Vaimberg, Katherine F Donovan, Ian Smith, et al. 2016. “Optimized SgRNA Design to Maximize Activity and Minimize Off-Target Effects of CRISPR-Cas9.” Nature Biotechnology 34 (2): 184–91. 10.1038/nbt.3437.

Doolittle, Lynda K., Michael K. Rosen, and Shae B. Padrick. 2013. “Measurement and Analysis of In Vitro Actin Polymerization.” In Adhesion Protein Protocols, edited by Amanda S. Coutts, 1046:273–93. Totowa, NJ: Humana Press. 10.1007/978-1-62703-538-5_16.

Ellis, Jonathan D., Miriam Barrios-Rodiles, Recep Çolak, Manuel Irimia, TaeHyung Kim, John A. Calarco, Xinchen Wang, et al. 2012. “Tissue-Specific Alternative Splicing Remodels Protein-Protein Interaction Networks.” Molecular Cell 46 (6): 884–92. 10.1016/j.molcel.2012.05.037.

Feather-Schussler, Danielle N., and Tanya S. Ferguson. 2016. “A Battery of Motor Tests in a Neonatal Mouse Model of Cerebral Palsy.” Journal of Visualized Experiments, no. 117 (November): 53569. 10.3791/53569.

Feng, Dairong, and Jiuyong Xie. 2013. “Aberrant Splicing in Neurological Diseases: Aberrant Splicing in Neurological Diseases.” Wiley Interdisciplinary Reviews: RNA 4 (6): 631–49. 10.1002/wrna.1184.

Fernandes, Jacqueline, Ivan M. Lorenzo, Yaniré N. Andrade, Anna Garcia-Elias, Selma A. Serra, José M. Fernández-Fernández, and Miguel A. Valverde. 2008. “IP3 Sensitizes TRPV4 Channel to the Mechano- and Osmotransducing Messenger 5′-6′-Epoxyeicosatrienoic Acid.” Journal of Cell Biology 181 (1): 143–55. 10.1083/jcb.200712058.

Fritz, Rafael D., Michel Letzelter, Andreas Reimann, Katrin Martin, Ludovico Fusco, Laila Ritsma, Bas Ponsioen, et al. 2013. “A Versatile Toolkit to Produce Sensitive FRET Biosensors to Visualize Signaling in Time and Space.” Science Signaling 6 (285). 10.1126/scisignal.2004135.

Gao, Chan, and Ye-Guang Chen. 2010. “Dishevelled: The Hub of Wnt Signaling.” Cellular Signalling 22 (5): 717–27. 10.1016/j.cellsig.2009.11.021.

García-Frigola, Cristina, Maria Isabel Carreres, Celia Vegar, Carol Mason, and Eloísa Herrera. 2008. “Zic2 Promotes Axonal Divergence at the Optic Chiasm Midline by EphB1-Dependent and -Independent Mechanisms.” Development 135 (10): 1833–41. 10.1242/dev.020693.

Gonatopoulos-Pournatzis, Thomas, and Benjamin J Blencowe. 2020. “Microexons: At the Nexus of Nervous System Development, Behaviour and Autism Spectrum Disorder.” Current Opinion in Genetics & Development 65 (December): 22–33. 10.1016/j.gde.2020.03.007.

Gonatopoulos-Pournatzis, Thomas, Mingkun Wu, Ulrich Braunschweig, Jonathan Roth, Hong Han, Andrew J. Best, Bushra Raj, et al. 2018. “Genome-Wide CRISPR-Cas9 Interrogation of Splicing Networks Reveals a Mechanism for Recognition of Autism-Misregulated Neuronal Microexons.” Molecular Cell 72 (3): 510-524.e12. 10.1016/j.molcel.2018.10.008.

Habas, Raymond, Yoichi Kato, and Xi He. 2001. “Wnt/Frizzled Activation of Rho Regulates Vertebrate Gastrulation and Requires a Novel Formin Homology Protein Daam1.” Cell 107 (7): 843–54.

Higashi, Tomohito, Tomoyuki Ikeda, Ryutaro Shirakawa, Hirokazu Kondo, Mitsunori Kawato, Masahito Horiguchi, Tomohiko Okuda, et al. 2008. “Biochemical Characterization of the Rho GTPase-Regulated Actin Assembly by Diaphanous-Related Formins, MDia1 and Daam1, in Platelets.” Journal of Biological Chemistry 283 (13): 8746–55. 10.1074/jbc.M707839200.

Irimia, Manuel, Robert J. Weatheritt, Jonathan D. Ellis, Neelroop N. Parikshak, Thomas Gonatopoulos-Pournatzis, Mariana Babor, Mathieu Quesnel-Vallières, et al. 2014. “A Highly Conserved Program of Neuronal Microexons Is Misregulated in Autistic Brains.” Cell 159 (7): 1511–23. 10.1016/j.cell.2014.11.035.

Jaiswal, Richa, Dennis Breitsprecher, Agnieszka Collins, Ivan R. Corrêa, Ming-Qun Xu, and Bruce L. Goode. 2013. “The Formin Daam1 and Fascin Directly Collaborate to Promote Filopodia Formation.” Current Biology 23 (14): 1373–79. 10.1016/j.cub.2013.06.013.

Kawabata Galbraith, Kelly, Kazuto Fujishima, Hiroaki Mizuno, Sung-Jin Lee, Takeshi Uemura, Kenji Sakimura, Masayoshi Mishina, Naoki Watanabe, and Mineko Kengaku. 2018. “MTSS1 Regulation of Actin-Nucleating Formin DAAM1 in Dendritic Filopodia Determines Final Dendritic Configuration of Purkinje Cells.” Cell Reports 24 (1): 95–106.e9. 10.1016/j.celrep.2018.06.013.

Kühn, Sonja, and Matthias Geyer. 2014. “Formins as Effector Proteins of Rho GTPases.” Small GTPases 5 (3): e983876. 10.4161/sgtp.29513.

Lamprecht, Raphael. 2021. “Actin Cytoskeleton Role in the Maintenance of Neuronal Morphology and Long-Term Memory.” Cells 10 (7): 1795. 10.3390/cells10071795.

Leger, Marianne, Anne Quiedeville, Valentine Bouet, Benoît Haelewyn, Michel Boulouard, Pascale Schumann-Bard, and Thomas Freret. 2013. “Object Recognition Test in Mice.” Nature Protocols 8 (12): 2531–37. 10.1038/nprot.2013.155.

Love, Michael I, Wolfgang Huber, and Simon Anders. 2014. “Moderated Estimation of Fold Change and Dispersion for RNA-Seq Data with DESeq2.” Genome Biology 15 (12): 550. 10.1186/s13059-014-0550-8.

Lu, Jun, Wuyi Meng, Florence Poy, Sankar Maiti, Bruce L. Goode, and Michael J. Eck. 2007. “Structure of the FH2 Domain of Daam1: Implications for Formin Regulation of Actin Assembly.” Journal of Molecular Biology 369 (5): 1258–69. 10.1016/j.jmb.2007.04.002.

Lueptow, Lindsay M. 2017. “Novel Object Recognition Test for the Investigation of Learning and Memory in Mice.” Journal of Visualized Experiments, no. 126 (August): 55718. 10.3791/55718.

Luo, Weijun, and Cory Brouwer. 2013. “Pathview: An R/Bioconductor Package for Pathway-Based Data Integration and Visualization.” Bioinformatics 29 (14): 1830–31. 10.1093/bioinformatics/btt285.

Mattila, Pieta K, and Pekka Lappalainen. 2008. “Filopodia: Molecular Architecture and Cellular Functions,” 9.

Matusek, T., R. Gombos, A. Szecsenyi, N. Sanchez-Soriano, A. Czibula, C. Pataki, A. Gedai, A. Prokop, I. Rasko, and J. Mihaly. 2008. “Formin Proteins of the DAAM Subfamily Play a Role during Axon Growth.” Journal of Neuroscience 28 (49): 13310–19. 10.1523/JNEUROSCI.2727-08.2008.

McLeod, Faye, and Patricia C Salinas. 2018. “Wnt Proteins as Modulators of Synaptic Plasticity.” Current Opinion in Neurobiology 53 (December): 90–95. 10.1016/j.conb.2018.06.003.

Messaoudi, E., T. Kanhema, J. Soule, A. Tiron, G. Dagyte, B. da Silva, and C. R. Bramham. 2007. “Sustained Arc/Arg3.1 Synthesis Controls Long-Term Potentiation Consolidation through Regulation of Local Actin Polymerization in the Dentate Gyrus In Vivo.” Journal of Neuroscience 27 (39): 10445–55. 10.1523/JNEUROSCI.2883-07.2007.

Nakaya, Masa-aki, Kristibjorn Orri Gudmundsson, Yuko Komiya, Jonathan R. Keller, Raymond Habas, Terry P. Yamaguchi, and Rieko Ajima. 2020. “Placental Defects Lead to Embryonic Lethality in Mice Lacking the Formin and PCP Proteins Daam1 and Daam2.” Edited by Michael Schubert. PLOS ONE 15 (4): e0232025. 10.1371/journal.pone.0232025.

Okuno, Hiroyuki, Kaori Akashi, Yuichiro Ishii, Nan Yagishita-Kyo, Kanzo Suzuki, Mio Nonaka, Takashi Kawashima, et al. 2012. “Inverse Synaptic Tagging of Inactive Synapses via Dynamic Interaction of Arc/Arg3.1 with CaMKIIβ.” Cell 149 (4): 886–98. 10.1016/j.cell.2012.02.062.

Otomo, Takanori, Diana R Tomchick, Chinatsu Otomo, Sanjay C Panchal, Mischa Machius, and Michael K Rosen. 2005. “Structural Basis of Actin FIlament Nucleation and Processive Capping by a Formin Homology 2 Domain” 433: 7.

Pan, Qun, Ofer Shai, Leo J Lee, Brendan J Frey, and Benjamin J Blencowe. 2008. “Deep Surveying of Alternative Splicing Complexity in the Human Transcriptome by High-Throughput Sequencing.” Nature Genetics 40 (12): 1413–15. 10.1038/ng.259.

Papandréou, Marie-Jeanne, and Christophe Leterrier. 2018. “The Functional Architecture of Axonal Actin.” Molecular and Cellular Neuroscience 91 (September): 151–59. 10.1016/j.mcn.2018.05.003.

Parras, Alberto, Héctor Anta, María Santos-Galindo, Vivek Swarup, Ainara Elorza, José L. Nieto-González, Sara Picó, et al. 2018. “Autism-like Phenotype and Risk Gene MRNA Deadenylation by CPEB4 Mis-Splicing.” Nature 560 (7719): 441–46. 10.1038/s41586-018-0423-5.

Plath, Niels, Ora Ohana, Björn Dammermann, Mick L. Errington, Dietmar Schmitz, Christina Gross, Xiaosong Mao, et al. 2006. “Arc/Arg3.1 Is Essential for the Consolidation of Synaptic Plasticity and Memories.” Neuron 52 (3): 437–44. 10.1016/j.neuron.2006.08.024.

Polder, Gerrit, Huub Hovens, and Hans Zweers. n.d. “Measuring Shoot Length of Submerged Aquatic Plants Using Graph Analysis,” 30.

Pollard, Thomas D. 2016. “Actin and Actin-Binding Proteins.” Cold Spring Harbor Perspectives in Biology 8 (8): a018226. 10.1101/cshperspect.a018226.

Quesnel-Vallières, Mathieu, Zahra Dargaei, Manuel Irimia, Thomas Gonatopoulos-Pournatzis, Joanna Y. Ip, Mingkun Wu, Timothy Sterne-Weiler, et al. 2016. “Misregulation of an Activity-Dependent Splicing Network as a Common Mechanism Underlying Autism Spectrum Disorders.” Molecular Cell 64 (6): 1023–34. 10.1016/j.molcel.2016.11.033.

Raj, Bushra, Manuel Irimia, Ulrich Braunschweig, Timothy Sterne-Weiler, Dave O’Hanlon, Zhen-Yuan Lin, Ginny I. Chen, et al. 2014. “A Global Regulatory Mechanism for Activating an Exon Network Required for Neurogenesis.” Molecular Cell 56 (1): 90–103. 10.1016/j.molcel.2014.08.011.

Ran, F Ann, Patrick D Hsu, Jason Wright, Vineeta Agarwala, David A Scott, and Feng Zhang. 2013. “Genome Engineering Using the CRISPR-Cas9 System.” Nature Protocols 8 (11): 2281–2308. 10.1038/nprot.2013.143.

Rial Verde, Emiliano M., Jane Lee-Osbourne, Paul F. Worley, Roberto Malinow, and Hollis T. Cline. 2006. “Increased Expression of the Immediate-Early Gene Arc/Arg3.1 Reduces AMPA Receptor-Mediated Synaptic Transmission.” Neuron 52 (3): 461–74. 10.1016/j.neuron.2006.09.031.

Roper, Randall J., Charles R. Goodlett, María Martínez de Lagrán, and Mara Dierssen. 2020. “Behavioral Phenotyping for Down Syndrome in Mice.” Current Protocols in Mouse Biology 10 (3): e79. 10.1002/cpmo.79.

Ruhela, Rakesh K, Shringika Soni, Phulen Sarma, Ajay Prakash, and Bikash Medhi. 2019. “Negative Geotaxis: An Early Age Behavioral Hallmark to VPA Rat Model of Autism.” Annals of Neurosciences 26 (1): 25–31. 10.5214/ans.0972.7531.260106.

Sakuma, Tetsushi, Ayami Nishikawa, Satoshi Kume, Kazuaki Chayama, and Takashi Yamamoto. 2015. “Multiplex Genome Engineering in Human Cells Using All-in-One CRISPR/Cas9 Vector System.” Scientific Reports 4 (1). 10.1038/srep05400.

Schindelin, Johannes, Ignacio Arganda-Carreras, Erwin Frise, Verena Kaynig, Mark Longair, Tobias Pietzsch, Stephan Preibisch, et al. 2012. “Fiji: An Open-Source Platform for Biological-Image Analysis.” Nature Methods 9 (7): 676–82. 10.1038/nmeth.2019.

Schönichen, André, and Matthias Geyer. 2010. “Fifteen Formins for an Actin Filament: A Molecular View on the Regulation of Human Formins.” Biochimica et Biophysica Acta (BBA) - Molecular Cell Research 1803 (2): 152–63. 10.1016/j.bbamcr.2010.01.014.

Shen, Wenjuan, Michaela B. C. Kilander, Morgan S. Bridi, Jeannine A. Frei, Robert F. Niescier, Shiyong Huang, and Yu-Chih Lin. 2020. “Tomosyn Regulates the Small RhoA GTPase to Control the Dendritic Stability of Neurons and the Surface Expression of AMPA Receptors.” Journal of Neuroscience Research 98 (6): 1213–31. 10.1002/jnr.24608.

Soneson, Charlotte, Michael I. Love, and Mark D. Robinson. 2015. “Differential Analyses for RNA-Seq: Transcript-Level Estimates Improve Gene-Level Inferences.” F1000Research 4 (December): 1521. 10.12688/f1000research.7563.1.

Szikora, Szilárd, István Földi, Krisztina Tóth, Ede Migh, Andrea Vig, Beáta Bugyi, József Maléth, et al. 2017. “The Formin DAAM Is Required for Coordination of the Actin and Microtubule Cytoskeleton in Axonal Growth Cones.” Journal of Cell Science, January, jcs.203455. 10.1242/jcs.203455.

Tapial, Javier, Kevin C.H. Ha, Timothy Sterne-Weiler, André Gohr, Ulrich Braunschweig, Antonio Hermoso-Pulido, Mathieu Quesnel-Vallières, et al. 2017. “An Atlas of Alternative Splicing Profiles and Functional Associations Reveals New Regulatory Programs and Genes That Simultaneously Express Multiple Major Isoforms.” Genome Research 27 (10): 1759–68. 10.1101/gr.220962.117.

Tomás Pereira, Inês, and Rebecca D. Burwell. 2015. “Using the Spatial Learning Index to Evaluate Performance on the Water Maze.” Behavioral Neuroscience 129 (4): 533–39. 10.1037/bne0000078.

Torres-Méndez. 2022. “Parallel Evolution of a Splicing Program Controlling Neuronal Excitability in Flies and Mammals.” SCIENCE ADVANCES, 20.

Torres-Méndez, Antonio, Sophie Bonnal, Yamile Marquez, Jonathan Roth, Marta Iglesias, Jon Permanyer, Isabel Almudí, et al. 2019. “A Novel Protein Domain in an Ancestral Splicing Factor Drove the Evolution of Neural Microexons.” Nature Ecology & Evolution 3 (4): 691–701. 10.1038/s41559-019-0813-6.

Vorhees, Charles V, and Michael T Williams. 2006. “Morris Water Maze: Procedures for Assessing Spatial and Related Forms of Learning and Memory.” Nature Protocols 1 (2): 848–58. 10.1038/nprot.2006.116.

Walf, Alicia A, and Cheryl A Frye. 2007. “The Use of the Elevated plus Maze as an Assay of Anxiety-Related Behavior in Rodents.” Nature Protocols 2 (2): 322–28. 10.1038/nprot.2007.44.

Wang, Eric T., Rickard Sandberg, Shujun Luo, Irina Khrebtukova, Lu Zhang, Christine Mayr, Stephen F. Kingsmore, Gary P. Schroth, and Christopher B. Burge. 2008. “Alternative Isoform Regulation in Human Tissue Transcriptomes.” Nature 456 (7221): 470–76. 10.1038/nature07509.

Warburton, E.C., and M.W. Brown. 2015. “Neural Circuitry for Rat Recognition Memory.” Behavioural Brain Research 285 (May): 131–39. 10.1016/j.bbr.2014.09.050.

Xu, Yingwu, James B Moseley, Isabelle Sagot, Florence Poy, David Pellman, Bruce L Goode, and Michael J Eck. n.d. “Crystal Structures of a Formin Homology-2 Domain Reveal a Tethered Dimer Architecture,” 13.

Yamashita, Masami, Tomohito Higashi, Shiro Suetsugu, Yusuke Sato, Tomoyuki Ikeda, Ryutaro Shirakawa, Toru Kita, et al. 2007. “Crystal Structure of Human DAAM1 Formin Homology 2 Domain: DAAM1 FH2 Structure.” Genes to Cells 12 (11): 1255–65. 10.1111/j.1365-2443.2007.01132.x.

Zhang, Haorui, Youssif Ben Zablah, Haiwang Zhang, and Zhengping Jia. 2021. “Rho Signaling in Synaptic Plasticity, Memory, and Brain Disorders.” Frontiers in Cell and Developmental Biology 9 (October): 729076. 10.3389/fcell.2021.729076.

Zhang, Hongyu, and Clive R. Bramham. 2021. “Arc/Arg3.1 Function in Long-term Synaptic Plasticity: Emerging Mechanisms and Unresolved Issues.” European Journal of Neuroscience 54 (8): 6696–6712. 10.1111/ejn.14958.

